# Mechanistic studies of mycobacterial glycolipid biosynthesis by the mannosyltransferase PimE

**DOI:** 10.1101/2024.09.17.613550

**Authors:** Yaqi Liu, Chelsea M. Brown, Nuno Borges, Rodrigo N. Nobre, Satchal Erramilli, Meagan Belcher Dufrisne, Brian Kloss, Sabrina Giacometti, Ana M. Esteves, Cristina G. Timóteo, Piotr Tokarz, Rosemary Cater, Yasu S. Morita, Anthony A. Kossiakoff, Helena Santos, Phillip J. Stansfeld, Rie Nygaard, Filippo Mancia

## Abstract

Tuberculosis (TB), exceeded in mortality only by COVID-19 among global infectious diseases, is caused by *Mycobacterium tuberculosis* (Mtb). The pathogenicity of Mtb is largely attributed to its complex cell envelope, which includes a class of glycolipids called phosphatidyl-*myo*-inositol mannosides (PIMs), found uniquely in mycobacteria and its related corynebacterineae. These glycolipids maintain the integrity of the mycobacterial cell envelope, regulate its permeability, and mediate host-pathogen interactions. PIMs consist of a phosphatidyl-*myo*-inositol core decorated with one to six mannose residues and up to four acyl chains. The mannosyltransferase PimE catalyzes the transfer of the fifth PIM mannose residue from a polyprenyl phosphate-mannose (PPM) donor. This step in the biosynthesis of higher-order PIMs contributes to the proper assembly and function of the mycobacterial cell envelope; however, the structural basis for substrate recognition and the catalytic mechanism of PimE remain poorly understood. Here, we present the cryo-electron microscopy (cryo-EM) structures of PimE from *Mycobacterium abscessus* captured in its apo form and in a product-bound complex with the reaction product Ac_1_PIM5 and the by-product polyprenyl phosphate (PP), determined at 3.0 Å and 3.5 Å, respectively. The structures reveal the active site within a distinctive binding cavity that accommodates both donor and acceptor substrates/products. Within the cavity, we identified residues involved in substrate coordination and catalysis, which we confirmed through *in vitro* enzymatic assays and further validated by *in vivo* complementation experiments. Molecular dynamics simulations were applied to identify the access pathways and the dynamics involved in substrate binding. Integrating structural, biochemical, genetic, and computational experiments, our study provides comprehensive insights into how PimE functions, opening potential avenues for development of novel anti-TB therapeutics.

## Introduction

Tuberculosis (TB) is an ancient disease that has plagued humanity throughout the ages^1,2^. It remains a significant global health challenge, with 7.5 million new cases and 1.3 million deaths reported in 2022 alone^3^. The causative agent of TB is the pathogenic bacterium *Mycobacterium tuberculosis* (Mtb). What distinguishes these pathogens is their cell envelope, which is a complex, multilayered structure that is hard to bypass and plays a crucial role in their survival and virulence^4–6^. It consists of an inner membrane rich in phospholipids and glycolipids, surrounded by a cell wall core composed of peptidoglycan covalently linked to arabinogalactan^7–10^. The cell wall core is esterified with long-chain mycolic acids, which form the inner leaflet of a unique outer membrane structure often referred to as the mycomembrane^11–14^. The outer leaflet of this mycomembrane is populated with lipids and glycolipids, including phosphatidylinositol mannosides (PIMs), lipomannan (LM), lipoarabinomannan (LAM), trehalose monomycolates (TMM), and trehalose dimycolates (TDM), among others^15–18^.

Among mycobacterial glycolipids of the inner membrane, PIMs are the most abundant^19–21^. They are characterized by the presence of one to six mannose residues and up to four acyl chains attached to their phosphatidyl-*myo*-inositol (PI) anchor^19,22^. PIMs are important components of the mycobacterial cell envelope, contributing to its structural integrity, regulating permeability, and mediating host-pathogen interactions^17,18,23,24^. Moreover, some PIMs are precursors of two large lipoglycans abundant in the mycobacterial cell envelope^25^, lipomannan (LM) and lipoarabinomannan (LAM), which are hyperglycosylated derivatives characterized by a linear mannan core, with LAM additionally containing arabinose branches^22,25^.

The early steps of the PIMs biosynthetic pathway occur on the cytoplasmic side of the inner membrane and have been partially characterized both at the genetic and biochemical levels^19,20^. The mannosyltransferase PimA (Rv2610c) and the mannosyltransferase PimB′ (Rv2188c) transfer mannose residues from GDP-mannose, to the 2-OH and 6-OH positions of the *myo*-inositol moiety of PI, respectively, yielding PIM1 and PIM2^26–28^ (Fig. 1a). The acyltransferase PatA (Rv2611c) then uses palmitoyl-CoA as a donor to acylate the 6-OH position of the mannose ring of PIM1 and PIM2, resulting in AcPIM1 and AcPIM2, respectively. An additional acyl chain moiety can be added to the 3-OH of the *myo*-inositol of AcPIM2 to form Ac_2_PIM2, but the enzyme carrying out this reaction remains unknown^29^. These acylated intermediates are presumed to translocate to the outer leaflet of the membrane, a process that is likely to require a currently uncharacterized flippase. Further mannosylation of Ac_1/2_PIM2 to Ac_1/2_PIM3 has been attributed to PimC, although the gene is only present in some strains of Mtb, suggesting that there is another unidentified gene that mediates the reaction^30,31^. The subsequent mannosylation of Ac_1/2_PIM3 to Ac_1/2_PIM4 is again presumed to be catalyzed by an unidentified enzyme denoted as PimD (Fig. 1a)^30,32^. The transfer of the fifth mannose to form Ac_1/2_PIM5 is known to be catalyzed by PimE^31^. Finally, the enzyme responsible for the final step of converting Ac_1/2_PIM5 to Ac_1/2_PIM6 remains unknown (Fig. 1a). A putative enzyme previously named PimF was initially thought to be involved in high molecular weight PIMs^33^; however, subsequent research has shown that this enzyme, now renamed LosA, is involved in the biosynthesis of an unrelated family of glycosylated acyltrehalose lipooligosaccharides in *M. marinum*^34^.

**Fig. 1.**
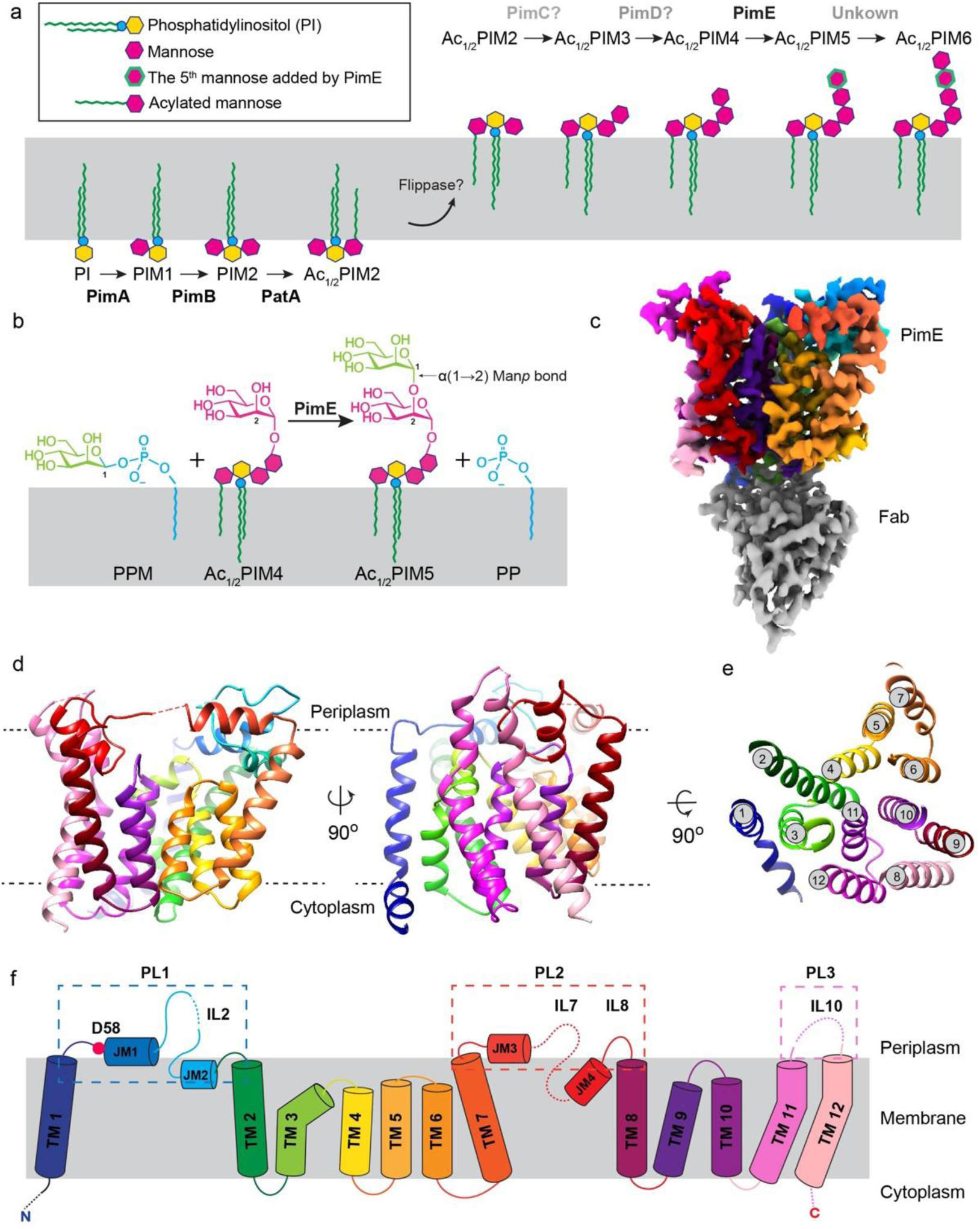
Biosynthetic Pathway and Structural Architecture of PimE. (a) Overview of PIM biosynthesis in mycobacteria, featuring the enzymatic reactions carried out by key mannosyltransferases. Enzyme names are indicated above the reaction arrows. The early steps occur on the cytoplasmic side of the plasma membrane, where PimA and PimB′ transfer mannose residues from GDP-mannose to the 2-OH and 6-OH positions of the inositol moiety of phosphatidylinositol (PI), yielding PIM1 and PIM2, respectively. PatA then acylates the 6-OH position of the mannose ring of PIM2, resulting in AcPIM2. An unknown acyltransferase can add an additional acyl chain to the 3-OH of the inositol of AcPIM2 to form Ac_2_PIM2. These acylated mannosylated intermediates are presumed to translocate to the outer leaflet of the membrane by an unknown flippase. PimC and PimD are thought to catalyze the subsequent tri- and tetra-mannosylation steps, leading to the formation of Ac_1_PIM4. PimE catalyzes the transfer of the fifth mannose residue from polyprenyl monophospho-β-D-mannose (PPM) toAc_1_PIM4, forming Ac_1_PIM5. The enzyme responsible for the conversion of Ac_1_PIM5 to AcPIM6 remains unknown. (b) Mannosyl transfer reaction catalyzed by PimE, showing the formation of an α(1→2) glycosidic bond between PPM and Ac_1_PIM4, yielding Ac_1_PIM5 and PP. Only tri-acylated PIMs (Ac_1_PIM4 and Ac_1_PIM5) are shown for the sake of simplicity. (c) Cryo-EM map of *Ma*PimE in complex with Fab-E6. PimE is depicted in rainbow, while the Fab-E6 is shown in grey. (d) PimE shown as ribbon colored in rainbow as in Fig. 1f. (e) TM helix arrangement of PimE is depicted as a cross-section colored in rainbow as in Fig. 1f. (f) Topological diagram of PimE showing the arrangement of TM segments and extracellular domains. The key catalytic residue D58 is shown as a red dot.

PimE, the focus of our studies, is a polyprenyl-monophospho-β-D-mannose (PPM)-dependent mannosyltransferase that catalyzes the transfer of the fifth mannose from a PPM donor to the Ac_1_PIM4 acceptor substrate, leading to the formation of Ac_1_PIM5 via an α(1→2) glycosidic bond (Fig. 1a-b)^31,35,36^. PPM is synthesized by PPM synthase (Ppm1) utilizing GDP-mannose and polyprenyl phosphate (PP) as precursors^37^. The length of the polyprenyl chain can vary, with the two most common forms being decaprenol (C50) and nonaprenol (C35)^36,38^. Previous studies have investigated the functional importance of specific residues in PimE, and D58 in *M. smegmatis* PimE, was found to be essential for enzymatic activity^31^. The genetic ablation of PimE leads to the accumulation of Ac_1/2_PIM4 and a deficiency in the synthesis of Ac_1/2_PIM6, which has been shown to have significant consequences for the structural integrity of the mycobacterial cell envelope and plasma membrane^31,39^.

While these findings underscore the relevance of PimE in the biosynthesis of PIMs, the structural basis for substrate recognition and catalysis by this enzyme remains elusive. In this study, we used single particle cryo-electron microscopy (cryo-EM) to determine high-resolution structures of PimE in the apo form and in a complex bound with its mannosylated product Ac_1_PIM5 and by-product PP at 3.0 Å and 3.5 Å resolution, respectively. Our structural analysis, complemented by molecular dynamics (MD) simulations and *in vitro* and *in vivo* functional assays, provides insights into the catalytic mechanism of PimE and the structural basis for substrate recognition.

## Results

### Structure determination of apo PimE

To identify a suitable candidate for structural studies, we screened PimE orthologs from 15 mycobacterial species, assessing their expression levels in *E. coli*, and solubility and stability in detergent. PimE from *Mycobacterium abscessus* (*Ma*PimE) exhibited the most promising properties and was therefore selected for further characterization. To confirm that *Ma*PimE retained enzymatic activity, we expressed it in *E. coli*, and incubated the isolated membrane fraction with its two substrates, Ac_1_PIM4 and PPM (see Materials and Methods for substrate preparation). The reaction products were analyzed by thin-layer chromatography (TLC), demonstrating the catalytic activity of recombinantly-expressed *Ma*PimE (Extended Data Fig. 1a).

For structure determination, *Ma*PimE was purified in n-Dodecyl-B-D-Maltoside (DDM) and reconstituted into lipid-filled nanodiscs (Extended Data Fig. 1c-d). *Ma*PimE has a molecular weight of 46 kDa, which poses a challenge for structure determination by cryo-EM^40,41^. To overcome this hurdle, we screened a synthetic phage display library to identify recombinant antigen-binding fragments (Fabs) that could specifically bind to *Ma*PimE, to increase the size of the particles and add extra-membrane feature to facilitate their alignment for cryo-EM data processing^40,42,43^. We evaluated eight Fab candidates for their ability to form stable complexes with *Ma*PimE, and we selected Fab-E6 due to its high binding affinity (Extended Data Fig. 1e-f).

We collected 7,469 micrographs of nanodisc-reconstituted apo *Ma*PimE in complex with Fab-E6 on a Titan Krios transmission microscope equipped with a K3 direct electron detector. After iterative 2D classification, we obtained high-quality 2D class averages showing bound Fab and secondary structural features within the nanodisc-embedded transmembrane (TM) domain. After further particle sorting and 3D refinement, we obtained a map with an overall resolution of 3.0 Å (Fig. 1c and Extended Data Fig. 2a-c).

### Structure of PimE

We were able to construct an almost complete atomic model of PimE with the exception of 20 residues at the N-terminus and 10 residues at the C-terminus, and three disordered loop regions (80-87, 246-250, 369-376) (Fig. 1d-f and Extended Data Fig. 3a). PimE contains twelve TM helices with both the N- and C-termini located on the cytoplasmic side of the membrane. These TM helices vary in length, ranging from 11 to 27 amino acids, with TM helix 12 being the longest (27 amino acids) and TM helix 6 the shortest (11 amino acids), spanning approximately two-thirds across the membrane (Fig. 1d-f, and Fig. 2a). The TM helices are interconnected by five short cytoplasmic loops (CL1-CL5), three periplasmic loops (PL1-PL3), three membrane-embedded loops and four juxtamembrane (JM) helices. Additionally, there are smaller connecting segments within these loops, particularly between the JM and the TM helices, designated interconnecting loops (IL). PL1, located between TM helices 1 and 2, contains two JM helices, JM1 and JM2, connected by a loop (IL2). JM1 is amphipathic and lies at the interface between the periplasmic space and the TM domain, parallel to the membrane plane (Fig. 1d-e and Fig. 2a). JM2 is positioned right below the interface, almost parallel to the membrane plane, and buried inside the membrane. PL2, which connects TM helices 7 and 8, also features two JM helices, JM3 and JM4, connected by a loop (IL7) (Fig. 1d-f). JM3 is situated at the periplasmic-membrane interface, while JM4 adopts an unusual orientation, being buried inside the membrane at an angle of approximately 60° relative to the plane of the membrane.

**Fig. 2.**
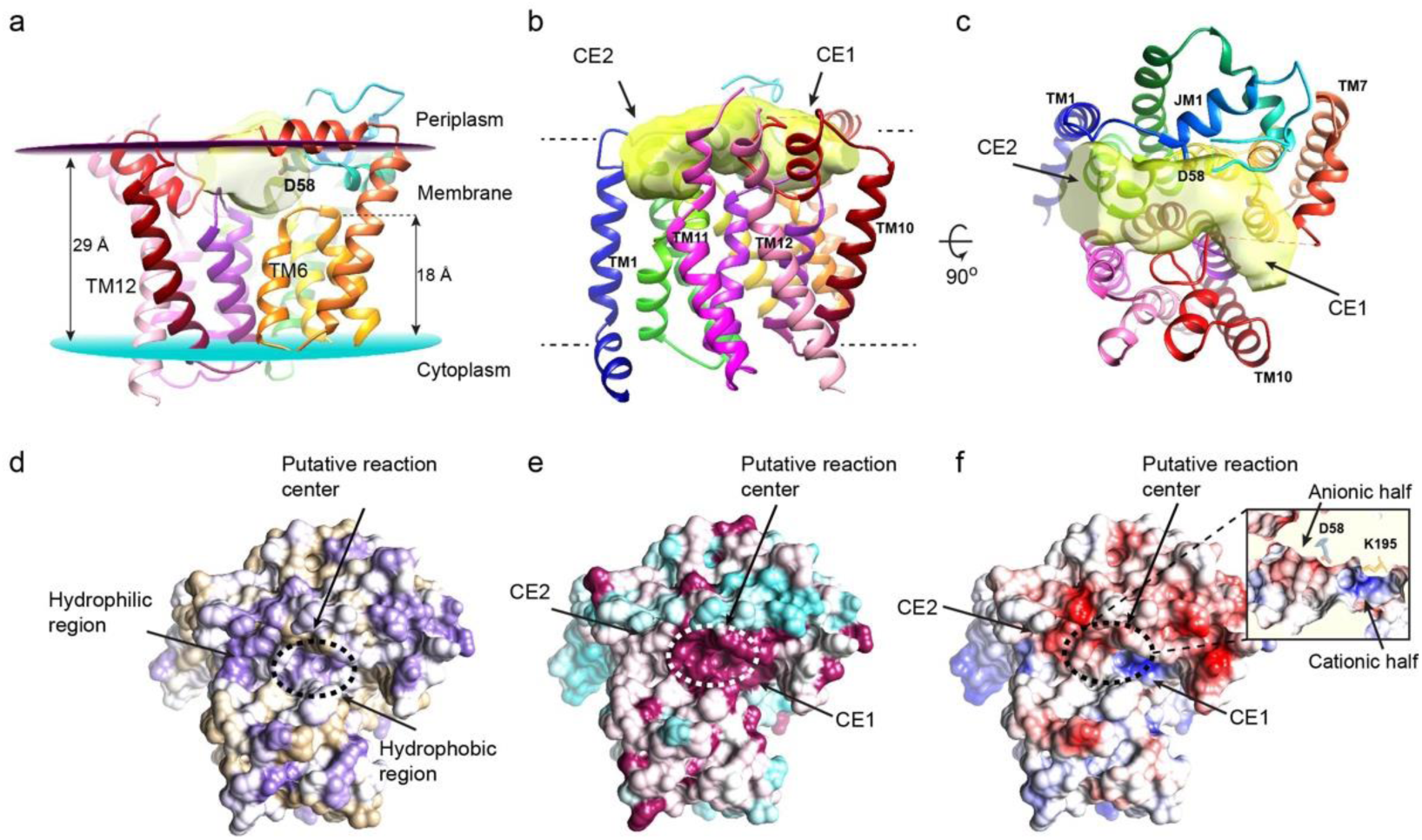
The putative substrate binding cavity of PimE. (a) The putative active site cavity within PimE is colored in semi-transparent yellow, showing its elongated, cashew-like shape and orientation relative to the membrane plane. (b-c) PimE viewed parallel (b) and perpendicular (c) to the membrane plane, with the elongated, cashew-shaped cavity shown in yellow. The cavity is oriented almost parallel to the membrane plane, with its two ends (Cavity End 1, CE1, and Cavity End 2, CE2) curving slightly towards the membrane. The cavity is surrounded by periplasmic loops (PL1 to PL3) and the connecting loops between TM helices 3 and 4 and TM helices 9 and 10. (d-f) PimE rendered in surface representation colored by hydrophobicity on a purple (very hydrophilic) to tan (very hydrophobic) scale (d), by electrostatic potential (e) on a range of ±5 kBT/e, and by conservation on a cyan (low conservation) to magenta (absolute conservation) scale (f). The putative reaction center at the central part of the cavity is marked with dotted circle.

The variable domain of the Fab was well resolved, allowing for reliable modeling. In contrast, the constant domain of the Fab was not included in the final model, due to the lack of well-defined density (Fig. 1f and Extended Data Fig. 3a). The cryo-EM map reveals well-resolved density at the interface between the Fab-E6 heavy chain and the cytoplasmic loops of *Ma*PimE (Extended Data Fig. 1g). Only the heavy chain appears to contribute to the interaction interface, which is stabilized through hydrogen bonds between its complementarity-determining regions (CDRs) and the cytoplasmic loops that connect TM helices 2 and 3, TM helices 4 and 5, and TM helices 8 and 9 in PimE (Extended Data Fig. 1g).

### Structural homology of PimE

Using the structural homology DALI server^44^ we confirmed that PimE belongs to the GT-C superfamily of glycosyltransferases. This superfamily can be further classified into the GT-C_A_ and GT-C_B_ subclasses^45^. The enzymes with the highest structural homology to PimE include yeast glucosyltransferase ALG6 (PDB: 6SNH)^46^, bacterial arabinosyltransferase ArnT (PDB: 5F15)^47^, and bacterial oligosaccharyltransferase PglB^48^ (PDB: 5OGL) (Extended Data Fig. 4a-d), all belonging to the GT-C_A_ subclass^45^. These enzymes share a conserved structural motif comprising, for PimE, the first seven TM helices (TM helices 1-7) and JM1 and JM2. Notably, D58 in *Ma*PimE, which corresponds to the previously identified catalytic residue D58 in *M. smegmatis* PimE^31^ (Extended Data Fig. 4a-d), is located at the apex of JM1. This position is equivalent to the catalytic aspartate residues in ALG6, ArnT, and PglB, further supporting its role as the essential catalytic base (Extended Data Fig. 4a-d). The DALI search also identified mycobacterial arabinosyltransferase AftA (PDB: 8IF8)^49^ as a structural homolog to PimE. While not explicitly classified in earlier reviews due to its recent structural publication, AftA likely also belongs to the GT-C_A_ subclass. Interestingly, the search also revealed structural similarities to proteins outside the GT-C family, such as PIGU, a subunit of the human GPI transamidase complex^50^ (PDB: 7WLD) (Extended Data Fig. 4e-f).

### Putative Substrate Binding Cavity of PimE

We identified a prominent cavity within PimE, located at the interface between the TM domain and the periplasm. This elongated cavity adopts a cashew-like shape and is oriented almost parallel to the membrane plane with its two ends curving slightly towards the membrane (Fig. 2a-c). The cavity is surrounded by three periplasmic loops (PL1, 2 and 3) and the connecting loops IL7 and IL9 (Fig. 1f and Fig. 2a-c). For clarity, we refer to the two ends of the cavity as “Cavity End 1” (CE1) and “Cavity End 2” (CE2) (Fig. 2b-c). CE1 is defined by JM3 and TM helices 5, 6, and 9, while CE2 is defined by JM1 and JM2 (Fig. 2b-c).

Given its location and structural features, we hypothesized that this cavity serves as the substrate binding site of PimE. The central part of the cavity, situated near the membrane-periplasm interface, is largely hydrophilic and highly conserved (Fig. 2d-e) and contains the predicted catalytic residue D58 (Fig. 2c). In contrast, the region of the cavity proximal to CE1 is predominantly hydrophobic, while the region on the CE2 side is more hydrophilic (Fig. 2d and Extended Data Fig. 6b). Furthermore, the electrostatic surface of the cavity reveals an uneven charge distribution, with the central region divided into two halves, a cationic half pointing towards CE1 and an anionic one towards CE2 (Fig. 2f and Extended Data Fig. 6c). The positive charge in the cationic half is partly caused by K195 which is highly conserved (Fig. 2f, Extended Data Fig. 5 and Extended Data Fig. 6c), while the anionic nature of the other half is mainly due to the presence of D58 (Fig. 2a-c). These properties suggest that the central region of the cavity serves as the reaction center (Fig. 2d-f), bringing together the hydrophilic head groups of the donor and acceptor substrates (Extended Fig. 6). We further hypothesized that CE1 acts as the entry point for the donor substrate PPM, with the hydrophobic region proximal to CE1 accommodating the polyprenyl tail of PPM. Conversely, CE2 could serve as the entry point for the acceptor Ac_1/2_PIM4, with its extended hydrophilic sugar head positioned along the cavity from the center towards CE2 and the acyl chain of Ac_1/2_PIM4 extending towards the TM domain.

### The cryo-EM structure of reaction products-bound *Ma*PimE

To validate our hypotheses and gain a deeper mechanistic understanding of this enzyme, we set forth to determine the structure of substrate-bound PimE. To this end, we added Ac_1_PIM4, isolated from the membranes of *M. smegmatis* mc^2^155 Δ*pimE*^51^, and PPM – enzymatically derived from *E. coli* BL21 (DE3) PLysS cells hosting a PPM synthase gene from *M. tuberculosis* H37Rv^52^– during the nanodisc incorporation stage of *Ma*PimE purification. We added Fab-E6 to this complex and determined its cryo-EM structure to 3.5 Å resolution following a similar flowchart to the one utilized for the apo structure (Extended Data Fig. 2d-f).

The resulting structure revealed an overall topology comparable to the apo structure (Fig. 3a-b). However, we observed two prominent and interpretable densities in the central cavity. One, close to CE1, was consistent with the phosphate group and the first three prenyl units of PPM. The absence of any density where we could fit the mannose moiety led us to interpret it as the PP portion of the PPM donor (Fig. 3 and Extended Data Fig. 7c). On the opposite side of the cavity, near CE2, the second density shows key characteristics of Ac_1_PIM5, and we could fit the inositol ring, five mannose residues, the phosphate group, and the first two carbons of the glycerol backbone directly connected to the phosphate group into this density (Fig. 3c-h and Extended Data Fig. 7c). However, the third carbon of the glycerol backbone and the acyl chains extending from it could not be traced reliably in the density map. Similarly, the acyl chain bound to the 6-OH position of the mannose residue linked to the 2-OH of the inositol remained elusive beyond the carbonyl group directly attached to the mannose (Fig. 3). The presence of PP and Ac_1_PIM5 in the structure, the products of the PimE reaction, despite having added PPM and Ac_1_PIM4, suggests that the mannosyl transfer reaction has occurred.

**Fig. 3.**
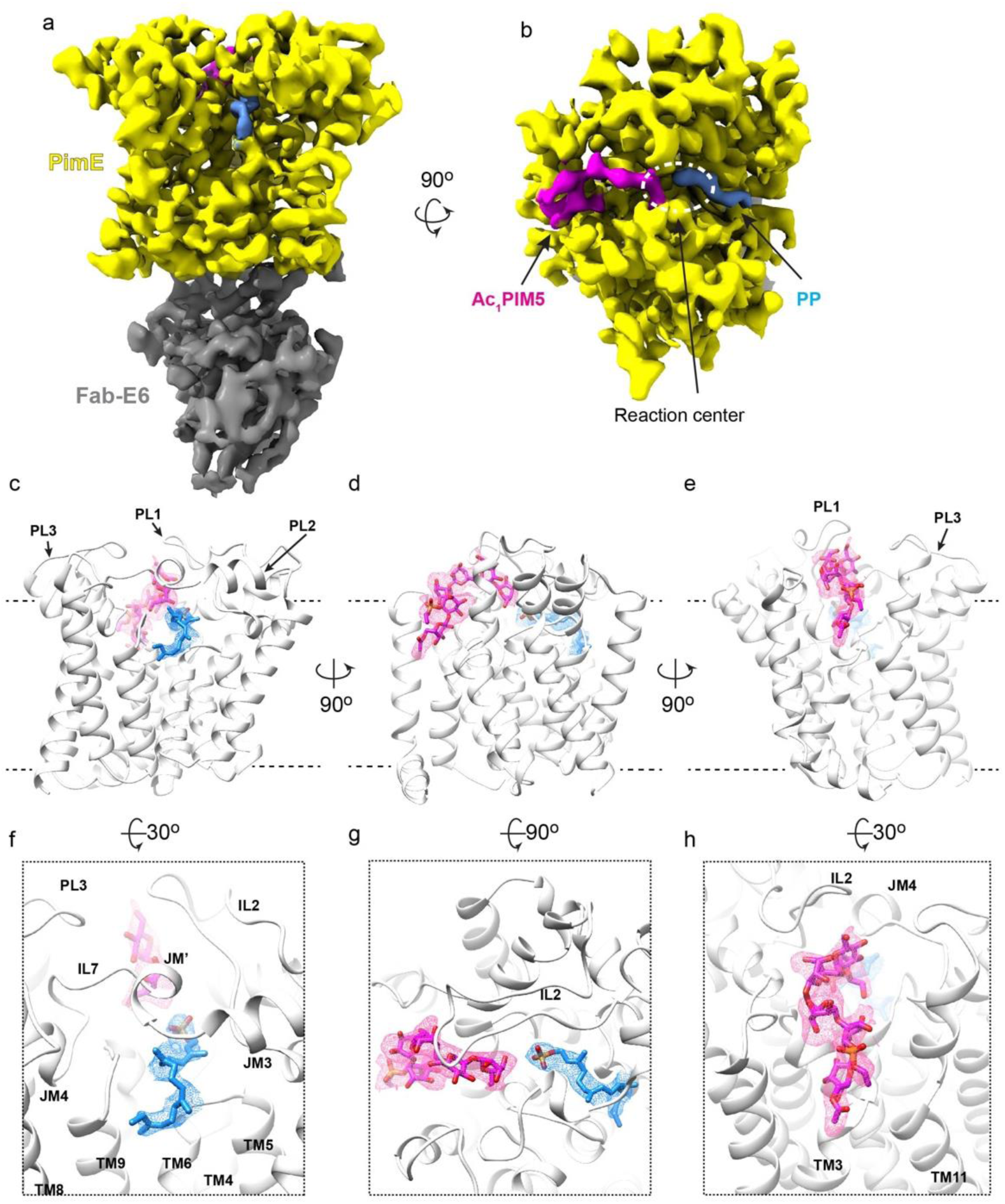
The cryo-EM structure of substrate-bound *Ma*PimE. (a-b) Cryo-EM density for the product-bound complex of *Ma*PimE with Ac_1_PIM5 (product) and PP (by-product), viewed parallel to the membrane plane. Fab-E6 binds to the same cytoplasmic domain of *Ma*PimE as observed in the apo structure. (c-e) Ribbon representation for the product-bound structure of PimE (white) bound with PP (blue) and Ac_1_PIM5 (magenta) shown in different orientations. PP and Ac_1_PIM5 adopt a curved shape, with their polar head groups projecting into the hydrophilic center of the substrate-binding cavity. (f-h) A focused view of the region where PP and Ac_1_PIM5 bind. PP and Ac_1_PIM5 adopt a curved shape. The polyprenyl chain of PP points towards the hydrophobic TM domain groove composed of TM helices 6 and 9, while Ac_1_PIM5 is surrounded by PL1, JM1, and JM4, with its acyl chains likely extending toward the TM region near TM helix 3.

Superimposing our two structures of PimE revealed subtle conformational rearrangements (Extended Data Fig. 6d). The TM domain of the product-bound structure exhibited minimal deviations from the apo structure, with a modest inward pivot rotation/translation of TM helix 7 being the most notable change. More noticeable rearrangements were observed in the PLs (Extended Data Fig. 6d). In PL1, the flexible region within IL2, connecting JM1 and JM2, which was previously disordered and unresolved in the apo structure, became traceable in the product-bound complex (Fig. 3c, Fig. 3f-h, and Extended Data Fig. 6d). Similarly, the extended loop (IL7) between JM3 and JM4 in PL2 was disordered in the apo form, but in the product-bound complex, IL7 became well-resolved, and an additional small helix (JM’) was formed (Fig. 3c and f, Extended Data Fig. 6d). This ordered IL7 appears to form an “arch” which the substrate threads its way under and into the active site cavity (Fig. 3c, 3f; Extended Data Fig. 4a, 6a). PL3, which connects TM helices 11 and 12, also transitioned from a partially disordered state in the apo form (Fig. 3c-f and Extended Data Fig. 6d) to a fully ordered and traceable conformation in the product-bound structure (Fig. 3c-f and Extended Data Fig. 6d).

Both PP and Ac_1_PIM5 adopt a curved shape, with their polar head groups projecting into the hydrophilic reaction center of the substrate-binding cavity (Fig. 3 and Extended Data Fig. 6a). The phosphate group of PP is located along the cationic half of the cavity (Extended Data Fig. 6c). At the same time, the mannose head of Ac_1_PIM5 is positioned along the anionic half (Extended Data Fig. 6c). Although only the first three prenyl units of the polyprenyl chain of PP could be resolved, the chain points towards the hydrophobic groove between TM helices 6 and 9 (Fig. 3f and Extended Data Fig. 7c). Based on the observed orientation and the hydrophobic nature of the groove, it is likely that the remaining prenyl units of the polyprenyl chain extend into this region (Fig. 3f). On the opposite side, Ac_1_PIM5 is surrounded by PL1, JM1, and JM4. While the acyl chains of Ac_1_PIM5 could not be fully traced in the map, their overall spatial arrangement suggests that they probably extend toward the membrane region near TM helix 3 (Fig. 3e and h).

### Structural and computational insights into ligand interactions

Our structure reveals a network of interactions between PimE and its ligands. At the center of the binding pocket, D58, located at the tip of JM1, forms a hydrogen bond with the C2 hydroxyl group of the fifth mannose of Ac_1_PIM5. Y62 forms a hydrogen bond with D58, potentially stabilizing its orientation within the active site. H321 engages with the fifth mannose moiety via a hydrogen bond. Y161 interacts with the glycosidic oxygen, linking the third and fourth mannose residues through a hydrogen bond. D55 coordinates the third mannose via a hydrogen bond with its hydroxyl group. W361 forms a hydrogen bond with the inositol moiety of Ac_1_PIM5, D160 coordinates with the inositol center through another hydrogen bond, and R43 establishes a hydrogen bond with the first mannose residue.

The phosphate group of PP is coordinated by three conserved residues H322, W319, and K195 (Fig. 4a). K195 forms a salt bridge with the phosphate, while H322 forms a hydrogen bond with one of the phosphate oxygens. W319 provides additional stabilization through a cation-π interaction with the positively charged K195 (Fig. 4a).

**Fig. 4.**
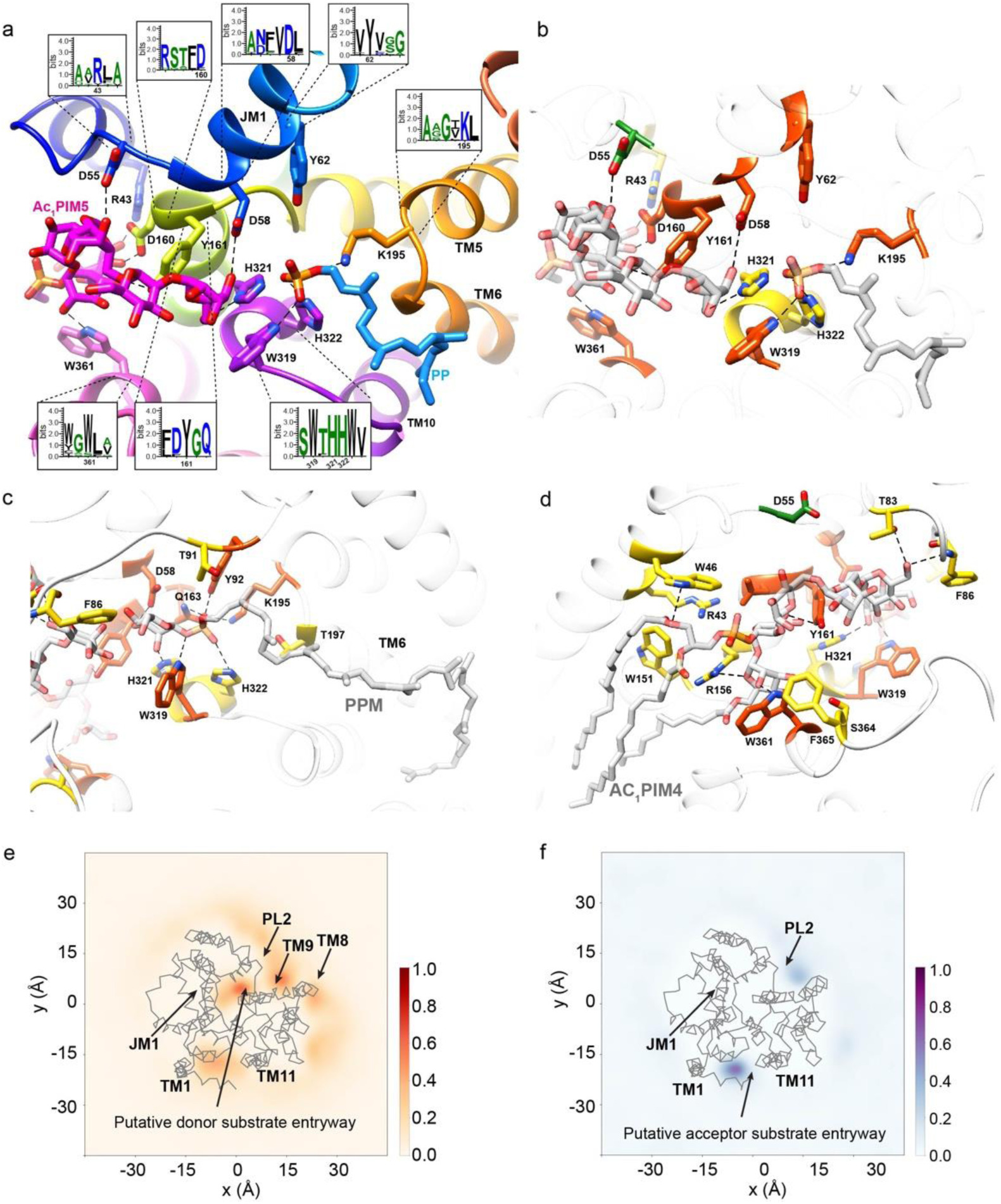
Structural insights into *Ma*PimE: Active site, substrate interactions, and functional residues. (a) Close-up view of the active site from the cryo-EM product-bound structure of PimE bound with products Ac_1_PIM5 and by-product PP. Key residues are shown as sticks. Insets: Sequence logos highlighting the conservation of active site residues involved in interactions with PP or Ac_1_PIM5. (b) Same view as (a), with residues colored according to mutational effects: red-orange for complete activity loss, yellow for reduced activity, and green for no change in activity. (c-d) RoseTTAFold docked models of PimE with full-length donor PPM and acceptor Ac_1_PIM4. Residues are colored as in (b) based on mutational effects. Hydrogen bonds are shown as black dotted lines in panels (a-d). (e-f) Density of PPM (e) and Ac_1_PIM4 (f) in CG-MD simulations with respect to PimE, with the protein backbone shown as gray lines. Regions of darker color show higher density by the lipid. Key areas of the protein are highlighted.

To complement our structural data and explore interactions with full-length substrates, we employed a combination of molecular docking and RoseTTAFold^54^. These computational approaches allowed us to model the full-length substrates and explore interactions not fully resolved in our cryo-EM structure.

The RoseTTAFold models confirm that H322, W319, and K195 coordinate the phosphate group of PPM, consistent with our observations from the cryo-EM structure. The models also show that Y92 and Q163 are involved in this coordination network through hydrogen bonding with the phosphate group (Fig. 4c).

For the donor substrate PPM, our computational studies indicate that H321 interacts with its mannose head group (Fig. 4c), complementing its observed interaction with the fifth sugar moiety of Ac_1_PIM5 in our cryo-EM structure (Fig. 4a). The models also show that the decaprenyl chain of PPM fits well into the hydrophobic groove, with the unresolved prenyl units extending along TM6, consistent with our structural observations.

In respect to the acceptor substrate, the RoseTTAFold model provides insights into interactions not fully captured in the cryo-EM structure. It suggests that Y161 coordinates the inositol moiety, while T83 and F86 coordinate the fourth mannose (Fig. 4d). R156 is shown to coordinate the first mannose. W46 is predicted to coordinate the ester oxygen that connects the sn-2 acyl chain to the glycerol backbone of the phosphatidylinositol in Ac_1_PIM5 (Fig. 4d). Our models also indicate the presence of several residues in the vicinity of the substrate binding sites, including W151, F365, S364, T91 and T197, which may contribute to substrate recognition, orientation, or active site organization (Fig. 4 c-d).

To further explore the dynamics of the cavity and the potential entry points for the substrates, we performed coarse-grained molecular dynamics (CG-MD) simulations. The simulations showed that the hydrophobic region proximal to CE1 favors the binding of PPM, with its lipid tail extending towards the TM domain (Fig. 4e). In contrast, the more hydrophilic region proximal to CE2 preferentially accommodates Ac_1_PIM4, with its sugar head positioned along the cavity from the center towards CE2 (Fig. 4f). These findings align with our structural analysis and provide additional insights into the substrate binding and entry mechanisms.

In addition to the CG-MD studies, simulations at atomistic resolution were performed in triplicate for both the substrates and products of the transglycosylase reaction. The ligands were stable and bound throughout both sets of 500 ns simulations (Extended Data Fig. 8a), indicating a high affinity for these positions. This can also be seen by the defined contact areas (Extended Data Fig. 8b-d) for both substrates and products. The increased stability of regions in the protein with the addition of the ligands seen in the resolved structures are also observed in the movement of the protein (Extended Data Fig. 9a-b) with and without ligands. The regions between JM1 and JM2 and the region between JM4 and JM5 are more dynamic when the binding partners are not present in the simulations. The simulations can provide further evidence to the key residues involved, with the *p*K_a_ of residues D58 and K195 fluctuating in different conditions (Extended Data Fig. 9c), suggesting a possible role in catalysis.

### Functional characterization of structure-based PimE mutants

To validate our structural and computational results, we designed specific mutations and performed *in vivo* complementation assays by expressing wild-type (WT) and mutant versions of *Ma*PimE in *M. smegmatis* Δ*pimE* and analyzed their PIMs profiles using thin-layer chromatography (TLC). As expected, the D58A and D58N mutations resulted in a complete loss of activity (Fig. 4b-c and Extended Data Fig. 10a-b). Nearby, Y62 forms a hydrogen bond with D58, and again not surprisingly, the Y62A mutation also led to a complete loss of activity (Fig. 4b-c and Extended Data Fig. 10a-b). Mutations affecting residues involved in coordinating the phosphate group of PPM and/or PP showed varying effects. K195A, Y92A, Q163A, and W319A mutations resulted in a complete loss of activity. In contrast, H321A and H322A mutants showed reduced activity, evidenced by some accumulation of Ac_1_PIM4 and detectable, albeit diminished, production of AcPIM6 (Fig. 4b-c and Extended Data Fig. 10a-b).

We also mutated several residues interacting with the acceptor substrate. D160A led to drastically reduced activity, while D160N showed PIMs profiles comparable to the WT enzyme, suggesting that hydrogen bonding capability at this position is sufficient for function (Fig. 4b and Extended Data Fig. 10a-b). Alanine mutations of W361 and Y161 resulted in a complete loss of activity (Fig. 4b and Extended Data Fig. 10c-f), whereas D55A and D55N mutations did not affect activity, suggesting a non-essential role for this residue (Fig. 4b and d and Extended Data Fig. 10a-b). R43A, T83A, F86A, and R156A all showed accumulation of Ac_1_PIM4 yet detectable production of AcPIM6, indicating reduced but not abolished activity (Fig. 4d and Extended Data Fig. 10c-f).

Furthermore, mutations of residues in the vicinity of the substrate binding sites (W151, F365, S364, T91, and T197) all showed accumulation of Ac_1_PIM4 yet detectable production of AcPIM6, indicating again reduced but not abolished activity (Fig. 4c-d and Extended Data Fig. 10e-f). This suggests that they contribute to efficient catalysis without being essential for the reaction.

To complement the functional data described above, we performed *in vitro* enzymatic assays using membrane fractions isolated from *E. coli* expressing WT and mutant PimE. Here, purified Ac_1_PIM4 and enzymatically synthesized PP[U-^14^C]M were added to the membrane fractions, and the formation of ^14^C-Ac_1_PIM5 was monitored by TLC. Consistent with our *in vivo* results (Extended Data Fig. 10a), D58A, D58N, K195A, W319A, and Y62A mutations completely abolished activity (Extended Data Fig. 7d-e, and Extended Data Fig. 10g). H321A and H322A showed reduced but not abolished activity. D55A and D55N mutations did not significantly affect activity. The D160A mutation largely reduced PimE activity, while D160N only partially reduced it (Extended Data Fig. 7d-e, and Extended Data Fig. 10g).

### Mechanism of substrate entry and catalysis

Metal ions play crucial roles in the catalytic activity of many glycosyltransferases^53^. However, the metal dependency of enzymes involved in PIMs biosynthesis, including PimE, remain poorly characterized. To investigate the potential metal dependency of PimE, we conducted enzymatic assays in the presence and absence of metal chelators (EGTA and EDTA). Our results showed that *Ma*PimE activity does not appear to be affected by the presence of these chelators (Extended Data Fig. 1b), suggesting that *Ma*PimE does not require metal ions for activity.

This observation, together with our analysis of the products-bound structure and insights from the MD simulations, allows us to propose a catalytic mechanism for PimE. We suggest that the donor substrate PPM accesses the active site cavity through an arch formed by IL7 (Fig. 5a). Once in the active site, the polar head of PPM is positioned with its phosphate group coordinated by the positively charged side chains of K195 and H322 (Fig. 5a). This arrangement allows the polyprenyl chain of PPM to align along the TM region formed by TM helices 6 and 9. On the opposite side of the active site, the sugar head moiety of the acceptor substrate Ac_1_PIM4 is positioned near the putative catalytic base D58 with its acyl chains extending towards TM helix 3 (Fig. 5a).

**Fig. 5.**
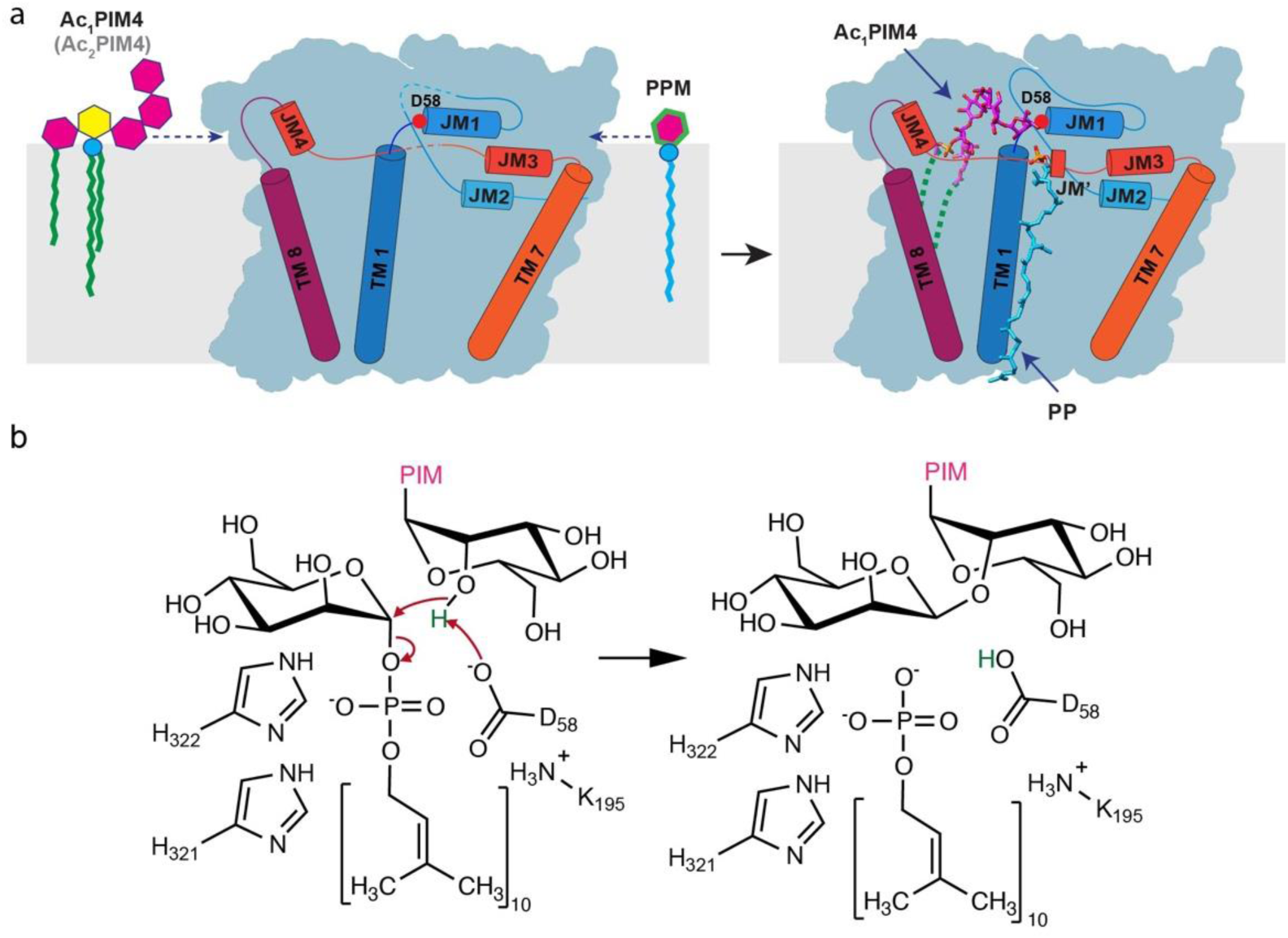
Proposed catalytic mechanism of PimE. (a) Schematic representation of the active site of PimE with bound substrates, showing the positioning of PPM and Ac_1_PIM4. D58, located at the tip of JM1, acts as the catalytic base. (b) Proposed catalytic mechanism, showing the role of D58 in deprotonating the 2-OH group of Ac_1_PIM4, initiating a nucleophilic attack on the anomeric carbon of PPM. This leads to the formation of an α(1→2) glycosidic bond between the mannose moiety of PPM and Ac_1_PIM4. The phosphate group of PPM is cleaved and stabilized by K195 and H322, forming the product Ac_1_PIM5 and the by-product PP. The tetra-acylated acceptor Ac_2_PIM4 and its corresponding glycosylated product Ac_2_PIM5 are not shown here for the sake of simplicity.

PimE utilizes PPM as the donor substrate and catalyzes the transfer of a mannosyl moiety from PPM to Ac_1_PIM4, forming an α(1→2) glycosidic bond and leading to the formation of Ac_1_PIM5. The stereochemical outcome classifies PimE as an inverting glycosyltransferase. Based on our structural observations and the general mechanism of inverting glycosyltransferases^56,57^, we propose that D58, located at the tip of JM1, acts as the catalytic base which deprotonates the 2-OH group of Ac_1_PIM4, initiating a nucleophilic attack on the anomeric carbon of PPM (Fig. 5b). This attack leads to the formation of an α(1→2) glycosidic bond between the mannose moiety of PPM and Ac_1_PIM4. As the glycosidic bond forms, the phosphate group of PPM is cleaved from the mannose moiety, becoming the leaving group. Our structural data and mutational studies suggest that K195 and H322 may play a role in stabilizing the negative charge of the phosphate group, facilitating its cleavage from PPM and the formation of Ac1PIM5 and the by-product PP (Fig. 5b).

## Discussion

PIMs are glycolipids present in abundance in the cell envelope of mycobacteria, that play critical roles in bacterial physiology and pathogenesis^18,25,54,55^. The biosynthesis of higher-order PIMs involves a series of mannosylation steps, with PimE catalyzing a late-stage reaction: the addition of the fifth mannose residue to form PIM5^31^. The high-resolution cryo-EM structures of *Ma*PimE in the apo and liganded form presented in this study provide insights into its architecture, ligand-binding properties, and catalytic mechanism. A distinctive feature of PimE is the presence of a cavity with a shape, and electrostatic and hydrophilic properties that enable it to accommodate both the tetramannoside sugar head of the acceptor substrate Ac_1/2_PIM4 and the donor substrate PPM. Notably, the cavity is flanked by distinct substrate entry points, with PPM accessing the active site through an arch-like structure formed by IL7, placing its long hydrophobic carbon chain along TM helix 6, and Ac_1/2_PIM4 entering from the opposite side, with its sugar head positioned near the catalytic base D58 (Fig. 5a).

PimE belongs to the GT-C_A_ subclass of glycosyltransferases, characterized by a conserved core module of seven TM helices. Structural comparisons of PimE with other GT-C_A_ glycosyltransferases, such as ALG6, ArnT, and PglB reveal a conserved mode of binding lipid-linked sugar donors^47,56,57^ (Extended Data Fig. 4a-d). In PimE, ALG6, and ArnT, the donor molecules access the active site through an arch-like structure formed by the loop bridging TM helices 7 and 8, while in PglB, this arch is formed by the loop between TM helices 9 and 10. Furthermore, the hydrophobic moiety of the donor substrate interacts with TM helix 6 in all these enzymes.

Our structural analysis also revealed similarities between PimE and PIGU (Extended Data Fig. 4f), a subunit of the human glycosylphosphatidylinositol (GPI) transamidase complex. While PimE is involved in the biosynthesis of PIMs, PIGU is responsible for attaching preformed GPI anchors to proteins in eukaryotic cells^50^. This structural resemblance is perhaps relevant given that PIMs and GPI anchors share as a common structural feature a phosphatidylinositol core decorated with mannose residues. Despite their distinct cellular contexts, the similarity in their structures suggests potential conservation of certain structural motifs involved in phosphatidylinositol-based glycolipid processing across different domains of life.

The GT-C_A_ architecture of PimE contrasts with the earlier enzymes in PIM biosynthesis, such as PimA and PimB′ (Fig. 1a). These enzymes are peripheral membrane-associated proteins belonging to the GT-B superfamily, characterized by two Rossmann fold-like domains^27,58^. The transition from these membrane-associated enzymes to the integral membrane protein PimE in the PIM biosynthesis pathway likely reflects the changing nature of the substrates as the synthesis progresses, particularly in respect to the donor substrate. While PimA and PimB′ utilize the soluble guanosine diphosphate (GDP)-mannose^27,28,58,59^, PimE uses the lipid-anchored PPM^64^. It has been speculated that PimE could also be responsible for adding the sixth mannose residue to Ac_1/2_PIM5^31^. We cannot confirm this hypothesis based on our structural and functional data, but it seems plausible, given that both reactions catalyze the formation of an α(1→2) glycosidic bond to incorporate the mannose residue.

Glycosyltransferases are broadly classified as either retaining or inverting enzymes depending on how they affect the stereochemistry at the anomeric center of the sugar they transfer^53,60^. PimE, like many other structurally characterized GT-C enzyme^55^, is classified as an inverting glycosyltransferase based on the resulting stereochemistry of the sugar product. The inverting glycosyltransferases are thought to operate through a single-step nucleophilic substitution mechanism, facilitated by an enzymatic general base catalyst, and can occur in either a metal dependent or independent way^60,61^. Our functional assays suggest that PimE activity is metal-independent, aligning with observations in structurally related enzymes like ALG6^46^ and AftA^62^ but contrasts with others such as PglB^48,57^. The metal dependency of earlier enzymes in the PIM biosynthesis pathway, including PimA and PimB′, remains unreported. In the context of this, based on our structural data, combined with computational studies and functional assays, we propose a catalytic mechanism for PimE. We suggest that D58 acts as the catalytic base, deprotonating the hydroxyl group of the acceptor substrate for nucleophilic attack on the anomeric carbon of the donor sugar. In metal-dependent glycosyltransferases, divalent cations often facilitate substrate binding, stabilize the transition state, and stabilize the leaving group^61,63^. In the absence of metal coordination, PimE appears to rely on the precise positioning of its catalytic residues and substrates within the active site. The orientation of D58 relative to the substrates is consistent with its proposed role as the catalytic base, while other conserved residues (such as K195 and H322) appear to be involved in substrate orientation, leaving group stabilization and possibly transition state stabilization. This proposed mechanism provides a starting point for understanding the stereochemical outcome of the reaction catalyzed by PimE.

The role of PimE in the biosynthesis of higher-order PIMs, vital for mycobacterial cell wall integrity and virulence, highlights its potential as a target for novel anti-tuberculosis therapeutics. The substrate binding cavity of PimE, with its distinct electrostatic properties and conserved residues involved in substrate recognition and catalysis, offers a potential binding site for small molecule inhibitor binding. Given the high conservation of PIM biosynthetic pathways across mycobacterial species, our findings are likely to extend to other clinically relevant mycobacteria, including *M. tuberculosis*.

In conclusion, our work provides a molecular framework for understanding PIM biosynthesis in mycobacteria. The structural insights and mechanistic details revealed here could serve as a foundation for structure-based drug design, potentially leading to novel anti-mycobacterial agents that target this critical cell envelope biosynthetic pathway.

## Methods

### Overexpression and purification of *Ma*PimE

WT *Ma*PimE was cloned into pNYCOMPSC23 plasmid^64^. The procedure for design and synthesis of pNYCOMPSC23 plasmid was described in detail in a previously published protocol^64^. The plasmids were transformed into BL21 (DE3) pLysS *E. coli* for protein expression. The transformed cells were grown at 37 °C to OD_600_ 0.6 - 0.8. The culture was then cooled to 22 °C and protein expression was induced by addition of 0.5 mM IPTG for 16 hours. Cells were harvested by centrifugation, the pellet was resuspended in lysis buffer containing 20 mM HEPES pH 7.5, 200 mM NaCl, 20 mM MgSO4, 10 mg/mL DNase I (Roche), 8 mg/mL RNase A (Roche), 1 mM tris(2-carboxyethyl) phosphine hydrochloride (TCEP), 1 mM PMSF, 1 tablet/1.5 L buffer EDTA-free cOmplete protease inhibitor cocktail (Roche). Cells were lysed by passing through a chilled Emulsiflex C3 homogenizer (Avestin) three times. The lysate was centrifuged at 3,000 x g in a Centrifuge 5810 R (Eppendorf) at 4°C for 5 min to remove cell debris and non-lysed cells. To isolate the cell membrane, the supernatant was ultracentrifuged in Type 45 Ti Rotor (Beckman Coulter) at 185,600 x g for 30 min. 4 L of culture, yielded ∼4-6 g of membrane pellet, which were resuspended to a final volume of 80-120 mL with lysis buffer, and homogenized using a handheld glass homogenizer (Kontes) on ice. The membrane fraction was stored at −80 °C until further use. The membrane fraction was thawed and solubilized by adding DDM to a final concentration of 1% (w/v) and was rotated gently at 4°C for 2 hours. Insoluble material was removed by ultracentrifugation at 185,600 x g in a Type 45 Ti Rotor at 4°C for 30 min. The supernatant was added to 3mL pre-equilibrated Ni^2+^-NTA resin (QIAGEN) in the presence of 40 mM imidazole and incubated with gentle rotation at 4 °C for 2 hours.

The resin was then washed with 10 column volumes (CV) of wash buffer containing 20mM HEPES pH 7.5, 200 mM NaCl, 65 mM Imidazole, 0.1% DDM and eluted with elution buffer containing 20 mM HEPES pH 7.5, 200mM NaCl, 300mM Imidazole, 0.03% DDM. The eluted protein was exchanged into a buffer containing 20 mM HEPES pH 7.5, 200mM NaCl, 0.03% DDM using a PD-10 desalting column (GE).

### Nanodisc reconstitution of PimE

The protein was then incorporated into lipid nanodisc (Bayburt and Sligar, 2010) with a molar ratio of 1:5:200 of membrane scaffold protein 1E3D1 (MSP1E3D1): 1-palmitoyl-2-oleoyl-glycero-3-phosphocholine (POPC) (Avanti), and incubated for 2 hours with gentle agitation at 4 °C. For the substrate bound structure, Ac_1_PIM4 and PPM were added to the reconstitution mixture at a molar ratio of 1:10:10 (PimE:Ac_1_PIM4:PPM). The POPC lipid was prepared by adding the solid extract to deionized water to a final concentration of 10 mM. The mixture was placed on ice and then gently sonicated with a tip sonicator (Fisher Scientific) to dissolve the lipids until it became semitransparent.

To initiate nanodisc reconstitution, 100 mg Biobeads (Bio-Rad) per mL of protein solution were added to the mixture and incubated by gentle rotation at 4 °C overnight. Biobeads were removed the next day by passing the reconstitution mixture through an Ultrafree centrifugal filter unit (Fisher) at 16,100 x g in a Centrifuge 5415 R (Eppendorf) at 4°C for 1 min. To remove free nanodisc, the reconstitution mixture was rebound to Ni^2+^-NTA resin with 20 mM imidazole for 2 hours at 4 °C. The resin was washed with 10 CV of wash buffer containing 20 mM HEPES pH 7.5, 200 mM NaCl and 40 mM imidazole, followed by 4 CV of elution buffer containing 20 mM HEPES pH 7.5, 200 mM NaCl and 300 mM imidazole. The eluted protein was subsequently purified by size-exclusion chromatography (SEC) with a Superdex 200 Increase 10/300 GL column in buffer containing 20 mM HEPES pH 7.5, 200 mM NaCl.

### Cryo-EM sample preparation

Fractions containing PimE incorporated into nanodiscs were pooled and incubated with Fab-E6 at 4 °C for 2 hours at a 1:3 molar ratio of protein to Fab. The protein mixture was further purified by SEC using a Superdex 200 Increase 10/300 GL column run in buffer containing 20 mM HEPES pH 7.5, 200 mM NaCl.

For the apo structure, peak fractions were pooled and concentrated to a final concentration of 5 mg/mL using a 50 kDa filter concentrator (Amicon). The sample was frozen using a Vitrobot (Thermo Fisher) by applying 3 μL of the sample to plasma cleaned (Gatan Solarus) 0.6/1-mm holey gold grid (Quantifoil UltrAuFoil). After a 30 second incubation, the grids were blotted using 595 filter paper (Ted Pella, Inc) for 8 seconds before immediately plunged into liquid ethane for vitrification. The plunger operated at 4 °C with greater than 90% humidity to minimize evaporation and sample degradation. For the substrate-bound structure, peak fractions were pooled and concentrated to a final concentration of 5 mg/mL. The grids were prepared the same way as for the apo sample, but with a blotting time of 9 seconds.

### Cryo-EM data collection

For the apo structure, images were recorded at the Columbia University Cryo-Electron Microscopy center on a Titan Krios electron microscope (FEI) equipped with an energy filter and a K3 direct electron detection filter camera (Gatan K3-BioQuantum) using a 0.87 Å pixel size. An energy filter slit width of 20 eV was used during the collection and was aligned automatically every hour using the Leginon software package (Suloway et al., 2005). Data collection was performed using a dose of approximately 58 e^−^ per Å^2^ across 50 frames (50 ms per frame) at a dose rate of around 16 e^−^ per pixel per second, using a set defocus range of −1.5 μm to −2.6 μm. A total of 15,198 micrographs were collected over a two-day session.

For the substrate-bound structure, images were recorded at the New York Structural Biology Center (NYSBC) on an FEI Titan Krios (NYSBC Krios 2) operating at 300 kV equipped with a spherical aberration corrector, an energy filter (Gatan GIF Quantum), and a post-GIF K2 Summit direct electron detector, using a 0.825 Å pixel size. An energy filter slit width of 20 eV was used during the collection and was aligned automatically every hour using the Leginon software package^65^. Data collection was performed using a dose of 50.3 e^−^ per Å^2^ across 24 frames (50 ms per frame) at a dose rate of around 41 e^−^ per pixel per second, using a set defocus range of −0.8 μm to −2.5 μm. A total of 21,900 micrographs were collected over a single session of four days.

### Identification of *Ma*PimE-specific Fab using phage display

*Ma*PimE was reconstituted into chemically biotinylated MSP1E3D1 as described above. Selection for Fabs was performed starting with Fab Library E^66,67^. Targets and the library were first diluted in selection buffer (20 mM HEPES, pH 7.4, 150 mM NaCl and 1% BSA). Five rounds of sorting were performed using a protocol adapted from published protocols^43,68^. In the first round, bio-panning was performed manually using 400 nM of *Ma*PimE, which was first immobilized onto magnetic beads and washed three times with selection buffer. The library was incubated for one hour with the immobilized target, beads were subsequently washed three times with selection buffer, and then beads were used to directly infect log-phase *E. coli* XL-1 Blue cells. Phages were amplified overnight in 2XYT media supplemented with ampicillin (100 µg/mL) and M13-K07 helper phage (10^9^ pfu/mL). To increase the stringency of selection pressure, four additional rounds of sorting were performed by stepwise reduction of the target concentration: 200 nM in the 2^nd^ round, 100 nM in the 3^rd^ round, and 50 nM in the 4^th^ and 5^th^ rounds. These rounds were performed semi-automatically using a KingFisher magnetic beads handler (Thermo Fisher Scientific). For each round, the amplified phage population from each preceding round was used as the input pool. Additionally, amplified phages were pre-cleared prior to each round using 100 µL of streptavidin paramagnetic particles, and 2.0 µM of empty MSP1E3D1 nanodiscs were used throughout the selection as competitors in solution. For rounds 2-5, prior to infection of log-phage cells, bound phage particles were eluted from streptavidin beads by a 15-minute incubation with 1% Fos-choline-12 (Anatrace).

### Single-point phage ELISA to validate Fab binding to *Ma*PimE

96-well plates (Nunc) were coated with 2 µg/mL Neutravidin and blocked with selection buffer. Colonies of *E. coli* XL-1 Blue cells harboring phagemids from the 4^th^ and 5^th^ rounds were used to inoculate 400 µL 2XYT media supplemented with 100 µg/mL ampicillin and 10^9^ pfu/mL M13-KO7 helper phage, and phages were subsequently amplified overnight in 96-well deep-well blocks with shaking at 280 rpm. Amplifications were cleared of cells with a centrifuge step and then diluted 10-fold into ELISA buffer (selection buffer with 2% BSA). All phages were tested against wells with immobilized biotinylated MSP1E3D1-reconstituted *Ma*PimE (30 nM), empty biotinylated-MSP1E3D1 nanodiscs (50 nM), or buffer alone to determine specific target binding. Phage ELISA was subsequently performed as previously described where the amount of bound phage was detected by colorimetric assay using an anti-M13 HRP-conjugated monoclonal antibody (GE Healthcare). Binders with high target and low non-specific signal were chosen for subsequent experiments.

### Fab cloning, expression and purification

Specific binders based on phage ELISA results were sequenced at the University of Chicago Comprehensive Cancer Center DNA Sequencing Facility and unique clones were then sub-cloned into the Fab expression vector RH2.2 (kind gift of S. Sidhu) using the In-Fusion Cloning kit (Takara). Successful cloning was verified by DNA sequencing. Fabs were then expressed and purified as previously described^69^. Following purification, Fab samples were verified for purity by SDS-PAGE and subsequently dialyzed overnight in 20 mM HEPES, pH 7.4, 150 mM NaCl.

### Assessment of Fab binding affinity to *Ma*PimE

To measure the apparent binding affinity, multi-point ELISAs using each purified Fab were performed in triplicate. Briefly, *M*aPimE (30 nM) or empty biotinylated-MSP1E3D1 nanodiscs (50 nM) were immobilized onto 96-well plates coated with Neutravidin (2 µg/mL). Fabs were diluted serially 3-fold into ELISA buffer using a starting concentration of 3 µM, and each dilution series was tested for binding to wells containing either *Ma*PimE, empty nanodiscs, or no target at all. The Fab ELISA was subsequently performed as previously described, where the amount of bound Fab was measured by a colorimetric assay using an HRP-conjugated anti-Fab monoclonal antibody (Jackson ImmunoResearch). Measured A_450_ values were plotted against the log Fab concentration, and EC_50_ values were determined in GraphPad Prism version 8.4.3 using a variable slope model assuming a sigmoidal dose-response.

### Single-particle cryo-EM data processing and map refinement

All data sets were corrected for beam-induced motion with Patch Motion Correction implemented in cryoSPARC v.2.15^70^ and the contrast transfer function (CTF) was estimated with Patch CTF. For the apo structure, 2.71 million particles were automatically picked using a blob-picker job and subjected to multiple rounds of 2D classification. Representative 2D classes of 80,420 particles clearly showing the Fab bound complex in the side views were selected as input for Topaz training. The resulting model was used to pick particles using Topaz Extract. Initially, 1.8 million particles were extracted in a box of 600 pixels and Fourier cropped to 150 pixels for initial cleanup.

Initially, 271,023 particles were extracted using a 320-pixel size box and binned four times. Multiple rounds of 2D classification were performed to clean up the particles using a batchsize of 400 per class and ‘‘Force Max over poses/shifts’’ turned off with 40 online-EM iterations. Classes with a total of 376,8 particles that displayed clear features of a Fab-bound nanodisc-embedded membrane protein were re-extracted using a 360-pixel box size without binning. *Ab initio* reconstruction was performed in cryoSPARC v.2.15 using two classes and a class similarity parameter of 0.1. From the *ab initio* reconstruction, one good class of 174,773 particles was subjected to heterogeneous refinement, and a final class of 145,477 particles was then subjected to non-uniform refinement and yielded a reconstruction with a resolution of 3.24 Å (FSC = 0.143). A subsequent local refinement was performed using a mask covering PimE and the variable region of the Fab and resulted in a density map at 3.02 Å resolution.

For the substrate-bound structure, 10.4 million particles were automatically picked using a blob-picker job and subjected to multiple rounds of 2D classification. Representative 2D classes of 103,632 particles clearly showing the Fab bound complex in the side views were selected as input for Topaz training. The resulting model was used to pick particles using Topaz Extract. Initially, 2.19 million particles were extracted in a box of 320 pixels and Fourier cropped to 80 pixels for initial cleanup. Multiple rounds of 2D classification were performed to clean up the particles using a batchsize of 400 per class and ‘‘Force Max over poses/shifts’’ turned off with 40 online-EM iterations. Classes with a total of 140,016 particles which displayed clear features of a Fab-bound nanodisc-embedded membrane protein were re-extracted using a 360-pixel box size without binning. *Ab initio* reconstruction was performed in cryoSPARC v.2.15 using two classes and a class similarity parameter of 0.1. One good class comprises of 78,648 particles were subjected to heterogeneous refinement and a final class comprised of 56,510 particles was subjected to non-uniform refinement and yielded a reconstruction with a resolution of 3.8 Å (FSC = 0.143). A subsequent local refinement was performed using a mask covering PimE and the variable region of the Fab and resulted in a density map at 3.46 Å resolution.

### Model building

Model building was performed in Coot^71^. Most of the Fab, except for the binding interface, was built based on a pre-existing model at high resolution (PDB ID 5UCB). The atomic model of apo PimE was built *de novo* from the globally sharpened 3.02 Å map using the Phenix Map to Model tool. Segments were manually joined, and residues were assigned based on the secondary structure prediction of Xtalpred^72^. Ramachandran outliers were manually fixed in Coot^71^ (version 0.9.6 EL). The structure was then refined using Phenix real space refine^76^ with secondary structure and Ramachandran restraints imposed. Models were validated using Molprobity within PHENIX 1.19.2^73,74^. For substrate-bound PimE, the model was built based on the apo structure, and the ligand files were generated in Coot (0.9.6 EL) and followed by iterative rounds of real-space refinement in PHENIX 1.19.2 and manual adjustment in Coot (0.9.6 EL).

### Expression and preparation of the PPM synthase

The second domain of the gene encoding a PPM synthase in *Mycobacterium tuberculosis* H37Rv, which starts with a methionine at position 594 and named ppm1_D2^37^ was cloned into the expression vector pET23a and introduced in *E. coli* BL21 (DE3) PLysS cells. Cells were grown in terrific broth medium supplemented with 100 μg/mL ampicillin and 34 μg/mL chloramphenicol at 37°C with agitation at 180 rpm until optical density at 600 nm reached 0.5-0.6. At this point, the culture was incubated at 4°C for 15 minutes. Protein expression was induced with 0.5 mM IPTG and growth was allowed to proceed at 16°C for 16 hours. Cells were harvested by centrifugation at 2,500 x g for 10 minutes at 4°C and resuspended in 25 mM Tris/HCl pH 7.6 supplemented with 10 mM MgCl_2_ and 5 mM β-mercaptoethanol. Cells were disrupted by sonication with cycles of 30 second of ice/disruption, repeated four times. Cell debris was removed by centrifugation at 3,000 x g for 30 minutes at 4°C and, after centrifugation, the supernatant was ultra-centrifuged at 84,000 x g for one hour at 4°C. The supernatant was discarded, and the membrane fraction resuspended gently in 25 mM Tris/HCl pH 7.6 supplemented with 10 mM MgCl_2_ and 5 mM β-mercaptoethanol. This membrane fraction was further used to produce PPM.

### Production of non-radiolabeled PPM

The membrane fraction of *E. coli* cells, harboring the PPM synthase gene, was incubated with 2.5 mM GDP-mannose, 0.1 mM ATP, 4 mM MgCl_2_, 50 mM Bicine buffer pH 8.0, and 1 mg of decaprenyl monophosphate (Larodan AB, Sweden) solubilized in CHAPS. The final concentration of CHAPS was 1% and the reaction mixture was at a final volume of 2.5 mL. The mixture was incubated for 1 hour at 37°C and the reaction quenched with the addition of 20.5 mL CHCl_3_/CH_3_OH/0.8M NaOH (10:10:3; v/v/v), and incubation at 50°C for 20 minutes. The mixture was allowed to cool down and purification of PPMs was carried out. Briefly, 1.75 mL of CHCl_3_ and 0.75 mL of H_2_0 were added and the mixture vortexed. The upper phase (aqueous phase) was discarded, and the bottom phase was washed three times with 2 mL of CHCl_3_/CH_3_OH/H_2_O (3:47:48; v/v/v). The lower phase was dried under a nitrogen stream and the residue resuspended in CHCl_3_/CH_3_OH/H_2_O (90:70:20; v/v/v). The PPM solution was loaded onto a silica gel 60 column (2.5 cm x 20 cm), prepared with 20 g of silica gel equilibrated with the same solvent system. Elution was carried out with the same solvent mixture: twenty fractions of 2 mL each were collected. The fractions were analyzed by High-performance thin-layer chromatography (HPTLC) by applying 10 μL of each. Visualization was performed, first by spraying with primulin and then with orcinol^51^. The fractions containing pure PPM were pooled and dried under a nitrogen stream. The residue was solubilized in 200 μL of CHCl_3_/CH_3_OH (2:1). The amount of PPM was estimated by total phosphorus quantification^75^.

### Production of radiolabeled PPM

The production of PP[U-^14^C]M was carried out as described for non-radiolabeled PPM, with the following modifications. The volume of the reaction mixture was decreased to 350 μL; the amount of decaprenyl monophosphate was 0.14 mg in 5% CHAPS. The final concentration of CHAPS in the reaction was 1%. Also, GDP-[U-^14^C] mannose (300 mCi/mmol) was added at a final concentration of 7.6 μM. The enzyme inactivation and purification of labeled PPM was carried out as described above, but the chromatographic step was excluded. The resulting PPM was resuspended in 200 μL of CHCl_3_/CH_3_OH (2:1, v/v) and quantification was done by scintillation counting. The absence of GDP-[U-^14^C] mannose was confirmed by TLC and the subsequent analysis was done by using the phosphor imaging plate and Fuji FLA-5100 imaging system.

### Determination of PimE activity in the membrane fraction

*E. coli* BL21 (DE3) pLysS cells harboring the *Mycobacterium abscessus* PimE gene (*Ma*PimE) were grown as described above. The cell pellet was resuspended in 25 mM HEPES pH 7.6 and disrupted by sonication using a tip sonicator (4 cycles of 30 seconds pulses with intervals of 30 seconds between pulses). Cell debris was removed by centrifugation (3,000 x g, 30 minutes, 4°C). The supernatant was ultracentrifuged (84,000 x g, 1 hour, 4°C), and the membrane fraction was gently resuspended in 25 mM HEPES pH 7.6. Total protein content was determined with the Pierce BCA protein assay kit. The *K_M_* and apparent *V_max_* of *Ma*PimE for PPM were evaluated in reaction mixtures (final volume, 30 μL), containing 50 mM Bicine buffer pH 8.0, 1 mM MgCl_2_, 1 mM Ac_1_PIM4, and different concentrations of PPM (50 μM to 1 mM; 2 μM PP[U-^14^C]M was included for each concentration examined). The *K_M_* and apparent *V_max_* for Ac_1_PIM4 was assessed in reaction mixtures (final volume, 30 μL), containing 50 mM of Bicine buffer pH 8.0, 1 mM MgCl_2_, 0.998 mM PPM, 2 μM PP[U-^14^C]M, and different concentrations of Ac_1_PIM4 (50 μM to 1 mM). Both substrates (PPM and Ac_1_PIM4) were solubilized in DDM, 1% (w/v) final concentration. The reaction mixtures were pre-incubated at 37 °C for 1 minute, and the reaction was initiated by addition of the membrane fraction (150 μg total protein) and incubated for a further 6 minutes; reactions were stopped by heating at 80 °C for 10 minutes. It was confirmed that the product formation is linear for up to 9 minutes of reaction time; furthermore, no Ac_1_PIM5 was produced during the enzyme inactivation step (Extended Data Fig. 7a-b).

Aliquots of the reaction mixtures (20 μL) were loaded onto a HPTLC plate and developed with chloroform/methanol/13 M ammonia/1 M ammonium acetate/water (180:140:9:9:23; v/v/v/v/v). The production of Ac_1_PIM5 was quantified by exposing, simultaneously, a HPTLC plate containing a known concentration of [U-^14^C]glucose (7.81 × 10^−4^ μCi). The HPTLC plate was exposed for 24 hours to a phosphor imaging plate; the image was obtained with a FujiFilm FLA-5100 imaging system. Quantification of the radiolabeled product, Ac_1_PIM5, was performed using the Fiji software^76^. *K_M_* and apparent *V_max_* values and respective standard deviations were calculated using the Origin software for nonlinear regression according to the Michaelis-Menten equation. All the reactions were performed in duplicate.

The two substrates, Ac_1_PIM4 and PPM, were produced in the lab since they are not commercially available. Ac_1_PIM4 was isolated from the membranes of *M. smegmatis* mc^2^155 Δ*pimE* as previously described^51^. PPM and PP[U-^14^C]M were obtained enzymatically from the membrane fraction of *E. coli* BL21 (DE3) PLysS cells, harboring the second domain of the gene encoding a PPM synthase from *M. tuberculosis* H37Rv. For a comprehensive description of the methods, see the Extended Data Material.

### *In vitro* test of activity of *Ma*PimE mutants

Point mutants of *Ma*PimE were generated using the QuikChange site-directed mutagenesis kit (Agilent). *E. coli* BL21 (DE3) cells harboring the WT and mutated genes were grown and membrane fractions were prepared as described above. Western blot analysis showed that all mutants have expression levels similar to the WT *Ma*PimE (Extended Data Fig. 10d). The activity of mutants and WT was evaluated under substrate saturation conditions (1 mM of each substrate) as described for the kinetic characterization of *Ma*PimE.

### Metal dependency assay for *Ma*PimE activity

*E. coli* membrane fractions expressing *Ma*PimE were prepared as described above. Reaction mixtures (30 μL) contained 50 mM Bicine buffer (pH 8.0), 1 mM MgCl_2_, membrane fraction (150 μg total protein), and substrates. Metal chelators (EGTA and EDTA) were added at 5 mM each. PPM and Ac_1_PIM4 were prepared as described below. Reactions were incubated at 37°C for 4 hours, then stopped by heating at 95°C for 5 minutes. Samples (3 μL) were spotted onto silica gel 60 HPTLC plates and developed using chloroform/methanol/13 M ammonia/1 M ammonium acetate/water (180:140:9:9:23, v/v/v/v/v) as the mobile phase. Glycolipids were visualized by orcinol staining.

### Mycobacterial strains and growth conditions

WT *M. smegmatis* mc²155^77^ and the Δ*pimE* mutant strain^31^ were used in this study. The Δ*pimE* strain was complemented with pYAMaPimE (encoding WT *M. abscessus* PimE) or its mutant derivatives. Strains were grown in Middlebrook 7H9 broth (Difco) supplemented with 10% DC (0.85% NaCl, 2% glucose), 0.05% Tween-80, and appropriate antibiotics (20 μg/mL kanamycin for Δ*pimE*; 20 μg/mL kanamycin plus 20 μg/mL streptomycin for complemented Δ*pimE* strains). Cultures were incubated at 37°C with shaking at 180 rpm until mid-log phase (OD_600_ 0.5-1.0).

### Construction of complementation plasmids and site-directed mutagenesis

The *M. abscessus pimE* gene was amplified by PCR and cloned into pYAB186^78^, replacing the *M. smegmatis pimE* gene to create the new plasmid pYAMaPimE. The pYAB186 vector contains both Flag and GFP tags^78^, which were retained in pYAMaPimE, resulting in Flag-GFP-tagged *Ma*PimE. Site-directed mutagenesis was performed using the QuikChange Lightning Kit (Agilent Technologies) following the manufacturer’s instructions. All constructs were verified by DNA sequencing.

### Transformation of *M. smegmatis*

Electrocompetent *M. smegmatis* Δ*pimE* cells were prepared and transformed as described previously^77^. Transformants were selected on Middlebrook 7H10 agar supplemented with 10% DC and appropriate antibiotics.

### Lipid extraction and HPTLC analysis

Lipid extraction and analysis were performed following an established protocol^82^. Briefly, bacterial cultures (50 OD_600_ units) were harvested by centrifugation at 3,214 x g for 10 minutes. Cell pellets were subjected to a three-step extraction process: first with 20 volumes of chloroform/methanol (2:1, v/v) for 1.5 hours, followed by two extractions with 10 volumes each of chloroform/methanol (2:1, v/v) and chloroform/methanol/water (1:2:0.8, v/v/v) for 1.5 hours each at room temperature. The combined organic extracts were dried under N_2_ stream and resuspended in water. The butanol phase was dried and resuspended in water-saturated butanol, and 3 μL (equivalent to 3 mg cell dry weight) were spotted on HPTLC silica gel 60 plates (Merck) and developed in chloroform/methanol/13 M NH_3_/1 M NH_4_Ac/water (180:140:9:9:23, v/v/v/v/v) for approximately 2 hours. Plates were dried and sprayed with orcinol spray reagent (0.1% orcinol in 20% H_2_SO_4_) and heated to visualize glycolipids.

### Western blot analysis

Wild-type, Δ*pimE*, and complemented *M. smegmatis* were cultured as described above. Log phase cells (OD_600_ = 0.5-1.0) were harvested by centrifugation and washed once with 50 mM HEPES buffer (pH 7.4). Cell pellets were resuspended in lysis buffer (25 mM HEPES pH 7.4, 2 mM EGTA) supplemented with EDTA-free cOmplete protease inhibitor cocktail (Roche). Cells were lysed by five passages through a chilled Emulsiflex C3 homogenizer (Avestin). The lysate was centrifuged at 10,000 x g for 15 minutes at 4°C to remove cell debris and unbroken cells. The supernatant was then ultracentrifuged in a Type 70 Ti Rotor (Beckman Coulter) at 126,000 x g for 30 min at 4°C to isolate the membrane fraction. The pellet was resuspended in lysis buffer and total protein content was determined using the Pierce BCA protein assay kit (Thermo Fisher Scientific). Equal amounts of protein from each sample were mixed with 4X Laemmli sample buffer containing 10% β-mercaptoethanol and separated on a 12% SDS-PAGE gel. Proteins were transferred to a PVDF membrane using the Trans-Blot Turbo Transfer System (Bio-Rad) following the manufacturer’s instructions.Flag-GFP-tagged PimE was detected using a monoclonal ANTI-FLAG M2-Peroxidase (HRP) antibody produced in mouse (1:5000 dilution; Sigma-Aldrich. Immunoreactive bands were visualized using the ECL Prime Western Blotting Detection Reagent (GE Healthcare) and imaged using Azure 600 ultimate Western blot imaging system (Azure Biosystems).

The membrane fractions of the wild-type *Ma*PimE and *Ma*PimE mutants expressed in *E. coli* were prepared as described above, and total protein content was determined using the Pierce BCA protein assay kit (Thermo Fisher Scientific). After electrophoresis on SDS-PAGE, proteins were transferred onto a PVDF membrane using the Trans-Blot Turbo Transfer System (Bio-Rad). His-tagged PimE mutants were directly detected with HisProbe-Horseradish Peroxidase (HRP) conjugate (1:5000 dilution; Thermo Fisher Scientific). His-tagged proteins were visualized using the ECL Prime Western Blotting Detection Reagent (GE Healthcare) in a ChemiDoc MP Imaging System (Bio-Rad).

### Coarse grained MD simulations

The CG parameters for PPM and Ac_1_PIM4 were generated based on previously published Martini 3 parameters^79–83^. For CG simulations, the protein was converted to the Martini 3 forcefield using martinize2^84^ including a 1000 kJ mol^−1^ nm^−2^ elastic network. PimE was then embedded into a PE:PG (80:20) membrane using the insane^85^ program, which was followed by solvation with martini water and neutralized with 150 M NaCl. For each of the substrate simulations, 4% of the upper leaflet of the membrane was this lipid. Minimization of the system was achieved using the steepest descents method. A timestep of 20 fs was used for production simulations, with five independent simulations of 5 µs for each substrate. The Parrinello-Rahman barostat^86^ was set at 1 bar and the velocity-rescaling thermostat^87^ was used at 310 K. All simulations were performed using Gromacs 2021.4^88,89^. For analysis, plumed^90^ was used in conjunction with matplotlib^91^ to generate the density plots.

### Atomistic simulations

Atomistic coordinates were generated from the end snapshots of the CG simulations using CG2AT2^92^. Molecular coordinates of both substrates and products were built using the coordinates from the ligand-bound cryo-EM structure. For the substrate state the transferred mannose was covalently added to the coordinates of the polyprenyl product and energy minimised to form coordinates for the PPM. RoseTTAFold All-atom was used to fold and dock both substrates and products for comparison. Atomistic molecular simulations were performed with the CHARMM36m forcefield, with parameters for Ac_1_PIM4 and Ac_1_PIM5 taken from CHARMM-GUI Membrane Builder^81,93^. Parameters for the PP and PPM were created using CHARMM-GUI and combined with previous parameters from our past studies. Energy minimizations of the systems were performed using the steepest descents method. A timestep of 2 fs was used for production simulations, with three independent simulations of 500 ns for substrate, product and apo states of PimE. The Parrinello-Rahman barostat^86^ was set at 1 bar and the velocity-rescaling thermostat^87^ was used at 310 K. All simulations were performed using Gromacs 2021.4^88,89^.

Analysis for the ligand contacts with the protein were performed with MDAnalysis^94^. The RMSF of the protein and RMSD of the ligands were measured using gmx tools. The *p*K_a_ of the residues were measured with PROPKA3^99^ and propkatraj (version 1.1.0)^100^. Plots were created with maplotlib^91^ and molecular visualization was performed with PyMOL.

## ACKNOWLEDGMENTS

This work was supported by R35GM132120 (to FM). PJS acknowledges the NIH (R01AI174416 (PI: M. Stephen Trent)), Wellcome (208361/Z/17/Z), MRC, BBSRC (BB/P01948X/1), and the Howard Dalton Centre for funding. CMB is supported by an MRC studentship (MR/N014294/1). PJS and CMB acknowledge Sulis at HPC Midlands+, which was funded by the EPSRC on grant EP/T022108/1, and the University of Warwick Scientific Computing Research Technology Platform for computational access. YSM acknowledges the NIH (R21AI168791) for funding. AAK acknowledges support from NIH grant R01GM117372. This project made use of time on ARCHER2 and JADE2 granted via the UK High-End Computing Consortium for Biomolecular Simulation, HECBioSim (http://www.hecbiosim.ac.uk), supported by EPSRC (grant no. EP/R029407/1). HS acknowledges Fundação para a Ciência e a Tecnologia (PTDC/BIA-BQM/31031/2017 (Lisboa-01-0145-FEDER-031031), MOSTMICRO-ITQB, UIDB/04612/2020 and UIDP/04612/2020). We thank Sara Rebelo for major technical assistance. We are especially grateful to Todd Lowary for his valuable input and insightful discussions that contributed to the improvement of this manuscript.

## AUTHOR CONTRIBUTIONS

Y.L. carried out protein expression, purification, cryo-EM sample preparation, data processing, model building, and structural refinement. B.K. conducted the high-throughput screening. M.B.D. and S.G. performed initial detergent screening to optimize purification conditions for *Ma*PimE. Y.L. performed site-directed mutagenesis for *Ma*PimE expressed in *E. coli*. R.N.N., N.B., A.M.E. and C.G.T. designed strategies and performed substrate production and purification as well as *in-vitro* functional assays under supervision of H.S. C.G.T. constructed the strain for PPM production and optimized growth conditions for PIMs production. Y.L. conducted HPTLC assays for metal-dependency testing of *Ma*PimE under the guidance of N.B. Y.L. performed molecular cloning for all *Ma*PimE constructs expressed in *M. smegmatis* and functional assays including mycobacterial complementation tests and HPTLC-based analysis of PIMs. Y.S.M. provided WT *M. smegmatis* mc²155, Δ*pimE* mutant strain, pYAB186 plasmid, and the competent cells for mycobacterial complementation tests, along with critical guidance for these experiments. C.M.B. and P.J.S. performed all molecular dynamics simulations and docking studies. S.E. and P.T. identified, characterized, and purified the Fabs. Y.L. drafted the manuscript with critical contributions from R.N., F.M., P.J.S., C.M.B., Y.S.M., N.B, A.M.E, H.S., S.E., and R.C., F.M., R.N., P.J.S., and H.S. supervised the project.

## DATA AND MATERIALS AVAILABILITY

## References

1. Daniel, T. M. The history of tuberculosis. Respir Med 100, 1862–1870 (2006).

2. Wirth, T. et al. Origin, spread and demography of the Mycobacterium tuberculosis complex. PLoS Pathog 4, (2008).

3. WHO. Report 20-23. January vol. t/malaria/ (2023).

4. Cirillo, J. D. & Kong, Y. Tuberculosis Host-Pathogen Interactions. Tuberculosis Host-Pathogen Interactions (Springer International Publishing, 2019). doi:10.1007/978-3-030-25381-3.

5. McNeil, M. R. & Brennan, P. J. Structure, function and biogenesis of the cell envelope of mycobacteria in relation to bacterial physiology, pathogenesis and drug resistance; some thoughts and possibilities arising from recent structural information. Res Microbiol 142, 451–463 (1991).

6. G. Lanéelle and M. Daffé. Myeobacterial cell wall and pathogenicity : a lipodologist’s view. Res Microbiol 142, 433–437 (1991).

7. Alderwick, L. J., Harrison, J., Lloyd, G. S. & Birch, H. L. The mycobacterial cell wall— peptidoglycan and arabinogalactan. Cold Spring Harb Perspect Med 5, 1–16 (2015).

8. Grzegorzewicz, A. E. et al. Assembling of the Mycobacterium tuberculosis cell wall core. Journal of Biological Chemistry 291, 18867–18879 (2016).

9. Kalscheuer, R. et al. The Mycobacterium tuberculosis capsule: A cell structure with key implications in pathogenesis. Biochemical Journal 476, 1995–2016 (2019).

10. Jackson, M., Crick, D. C. & Brennan, P. J. Phosphatidylinositol is an essential phospholipid of mycobacteria. Journal of Biological Chemistry 275, 30092–30099 (2000).

11. Marrakchi, H., Lanéelle, M. A. & Daffé, M. Mycolic acids: Structures, biosynthesis, and beyond. Chem Biol 21, 67–85 (2014).

12. Chiaradia, L. et al. Dissecting the mycobacterial cell envelope and defining the composition of the native mycomembrane. Sci Rep 7, 1–12 (2017).

13. Charles L. Dulberger, E. J. R. & C. C. B. The mycobacterial cell envelope — a moving target. Nat Rev Microbiol 47–59 (2020) doi:10.5040/9781501303746.ch-002.

14. Hoffmann, C., Leis, A., Niederweis, M., Plitzko, J. M. & Engelhardt, H. Disclosure of the mycobacterial outer membrane: Cryo-electron tomography and vitreous sections reveal the lipid bilayer structure. Proc Natl Acad Sci U S A 105, 3963–3967 (2008).

15. Jackson, M. The mycobacterial cell envelope-lipids. Cold Spring Harb Perspect Med 4, 1–22 (2014).

16. Lee, R. E., Brennan, P. J. & Besra, G. S. Mycobacterium tuberculosis cell envelope. Curr Top Microbiol Immunol 215, 1–27 (1996).

17. Fukuda, T. et al. Critical roles for lipomannan and lipoarabinomannan in cell wall integrity of mycobacteria and pathogenesis of tuberculosis. mBio 4, 8–10 (2013).

18. Angala, S. K., Belardinelli, J. M., Huc-Claustre, E., Wheat, W. H. & Jackson, M. The Cell Envelope Glycoconjugates of Mycobacterium Tuberculosis. Critical Reviews in Biochemistry and Molecular Biology vol. 49 (2014).

19. Guerin, M. E., Korduláková, J., Alzari, P. M., Brennan, P. J. & Jackson, M. Molecular Basis of Phosphatidyl-myo-inositol Mannoside Biosynthesis and Regulation in Mycobacteria. Journal of Biological Chemistry 285, 33577–33583 (2010).

20. Sancho-Vaello, E., Albesa-Jové, D., Rodrigo-Unzueta, A. & Guerin, M. E. Structural basis of phosphatidyl-myo-inositol mannosides biosynthesis in mycobacteria. Biochim Biophys Acta Mol Cell Biol Lipids 1862, 1355–1367 (2017).

21. Nigou, J., Gilleron, M. & Puzo, G. Lipoarabinomannans: From structure to biosynthesis. Biochimie 85, 153–166 (2003).

22. Angala, S. K., Belardinelli, J. M., Huc-Claustre, E., Wheat, W. H. & Jackson, M. The cell envelope glycoconjugates of Mycobacterium tuberculosis. Critical Reviews in Biochemistry and Molecular Biology vol. 49 361–399 Preprint at 10.3109/10409238.2014.925420 (2014).

23. Chatterjee, D. & Khoo, K. H. Mycobacterial lipoarabinomannan: An extraordinary lipoheteroglycan with profound physiological effects. Glycobiology 8, 113–120 (1998).

24. Briken, V., Porcelli, S. A., Besra, G. S. & Kremer, L. Mycobacterial lipoarabinomannan and related lipoglycans: From biogenesis to modulation of the immune response. Mol Microbiol 53, 391–403 (2004).

25. Källenius, G., Correia-Neves, M., Buteme, H., Hamasur, B. & Svenson, S. B. Lipoarabinomannan, and its related glycolipids, induce divergent and opposing immune responses to Mycobacterium tuberculosis depending on structural diversity and experimental variations. Tuberculosis 96, 120–130 (2016).

26. Korduláková, J. et al. Identification of the required acyltransferase step in the biosynthesis of the phosphatidylinositol mannosides of Mycobacterium species. Journal of Biological Chemistry 278, 36285–36295 (2003).

27. Guerin, M. E. et al. Molecular recognition and interfacial catalysis by the essential phosphatidylinositol mannosyltransferase PimA from mycobacteria. Journal of Biological Chemistry 282, 20705–20714 (2007).

28. Korduláková, J. et al. Definition of the first mannosylation step in phosphatidylinositol mannoside synthesis: PimA is essential for growth of mycobacteria. Journal of Biological Chemistry 277, 31335–31344 (2002).

29. Mishra, A. K. et al. Identification of a novel α(1→6) mannopyranosyltransferase MptB from Corynebacterium glutamicum by deletion of a conserved gene, NCgl1505, affords a lipomannan- and lipoarabinomannan-deficient mutant. Mol Microbiol 68, 1595–1613 (2008).

30. Kremer, L. et al. Characterization of a putative α-mannosyltransferase involved in phosphatidylinositol trimannoside biosynthesis in Mycobacterium tuberculosis. Biochemical Journal 363, 437–447 (2002).

31. Morita, Y. S. et al. PimE is a polyprenol-phosphate-mannose-dependent mannosyltransferase that transfers the fifth mannose of phosphatidylinositol mannoside in mycobacteria. Journal of Biological Chemistry 281, 25143–25155 (2006).

32. Berg, S., Kaur, D., Jackson, M. & Brennan, P. J. The glycosyltransferases of Mycobacterium tuberculosis - Roles in the synthesis of arabinogalactan, lipoarabinomannan, and other glycoconjugates. Glycobiology 17, (2007).

33. Alexander, D. C., Jones, J. R. W., Tan, T., Chen, J. M. & Liu, J. PimF, a Mannosyltransferase of Mycobacteria, Is Involved in the Biosynthesis of Phosphatidylinositol Mannosides and Lipoarabinomannan. Journal of Biological Chemistry 279, 18824–18833 (2004).

34. Burguière, A. et al. LosA, a key glycosyltransferase involved in the biosynthesis of a novel family of glycosylated acyltrehalose lipooligosaccharides from Mycobacterium marinum. Journal of Biological Chemistry 280, 42124–42133 (2005).

35. Kovacevic, S. et al. Identification of a novel protein with a role in lipoarabinomannan biosynthesis in mycobacteria. Journal of Biological Chemistry 281, 9011–9017 (2006).

36. Wolucka, B. A. & De Hoffmann, E. Isolation and characterization of the major form of polyprenyl-phospho-mannose from Mycobacterium smegmatis. Glycobiology 8, 955–962 (1998).

37. Gurcha, S. S. et al. Ppm1, a novel polyprenol monophosphomannose synthase from Mycobacterium tuberculosis. Biochemical Journal 365, 441–450 (2002).

38. K Takayama, H K Schnoes, E. J. S. Characterization of the alkali-stable mannophospholipids of Mycobacterium smegmatis. Biochim Biophys Acta Aug 23;316, 212–21 (1973).

39. Crellin, P. K. et al. Mutations in pimE restore lipoarabinomannan synthesis and growth in a Mycobacterium smegmatis lpqW mutant. J Bacteriol 190, 3690–3699 (2008).

40. Nygaard, R., Kim, J. & Mancia, F. Cryo-electron microscopy analysis of small membrane proteins. Curr Opin Struct Biol 64, 26–33 (2020).

41. de Oliveira, T. M., van Beek, L., Shilliday, F., Debreczeni, J. & Phillips, C. Cryo-EM: The Resolution Revolution and Drug Discovery. SLAS Discovery 26, 17–31 (2021).

42. Wentinck, K., Gogou, C. & Meijer, D. H. Putting on molecular weight: Enabling cryo-EM structure determination of sub-100-kDa proteins. Curr Res Struct Biol 4, 332–337 (2022).

43. Paduch, M. & Kossiakoff, A. A. Generating Conformation and Complex-Specific Synthetic Antibodies. 1575, 93–119 (2017).

44. Holm, L. & Laakso, L. M. Dali server update. Nucleic Acids Res 44, W351–W355 (2016).

45. Alexander, J. A. N. & Locher, K. P. Emerging structural insights into C-type glycosyltransferases. Curr Opin Struct Biol 79, 102547 (2023).

46. Bloch, J. S. et al. Structure and mechanism of the ER-based glucosyltransferase ALG6. Nature 579, 443–447 (2020).

47. Petrou, V. I. et al. Structural biology: Structures of aminoarabinose transferase ArnT suggest a molecular basis for lipid A glycosylation. Science (1979) 351, 608–612 (2016).

48. Napiórkowska, M., Boilevin, J., Darbre, T., Reymond, J. L. & Locher, K. P. Structure of bacterial oligosaccharyltransferase PglB bound to a reactive LLO and an inhibitory peptide. Sci Rep 8, 1–9 (2018).

49. Gong, Y. et al. Structure of the priming arabinosyltransferase AftA required for AG biosynthesis of Mycobacterium tuberculosis. Proc Natl Acad Sci U S A 120, 1–7 (2023).

50. Xu, Y. et al. Molecular insights into biogenesis of glycosylphosphatidylinositol anchor proteins. Nat Commun 13, 1–13 (2022).

51. Nobre, R. N. et al. Production and Purification of Phosphatidylinositol Mannosides from Mycobacterium smegmatis Biomass. Curr Protoc 2, 1–22 (2022).

52. Scherman, H. et al. Identification of a polyprenylphosphomannosyl synthase involved in the synthesis of mycobacterial mannosides. J Bacteriol 191, 6769–6772 (2009).

53. Moremen, K. W. & Haltiwanger, R. S. Emerging structural insights into glycosyltransferase-Mediated Synthesis of Glycans. Nat Chem Biol 15, 853–864 (2019).

54. Court, N. et al. Mycobacterial PIMs inhibit host inflammatory responses through CD14-dependent and CD14-independent mechanisms. PLoS One 6, (2011).

55. Ramon-Luing, L. A., Palacios, Y., Ruiz, A., Téllez-Navarrete, N. A. & Chavez-Galan, L. Virulence Factors of Mycobacterium tuberculosis as Modulators of Cell Death Mechanisms. Pathogens 12, (2023).

56. Bloch, J. S. et al. Structure and mechanism of the ER-based glucosyltransferase ALG6. Nature 579, 443–447 (2020).

57. Napiórkowska, M. et al. Molecular basis of lipid-linked oligosaccharide recognition and processing by bacterial oligosaccharyltransferase. Nat Struct Mol Biol 24, 1100–1106 (2017).

58. Batt, S. M. et al. Acceptor substrate discrimination in phosphatidyl-myo-inositol mannoside synthesis: Structural and mutational analysis of mannosyltransferase corynebacterium glutamicum PimB′. Journal of Biological Chemistry 285, 37741–37752 (2010).

59. Guerin, M. E. et al. New insights into the early steps of phosphatidylinositol mannoside biosynthesis in mycobacteria: PimB′ is an essential enzyme of Mycobacterium smegmatis. Journal of Biological Chemistry 284, 25687–25696 (2009).

60. Breton, C., Šnajdrová, L., Jeanneau, C., Koča, J. & Imberty, A. Structures and mechanisms of glycosyltransferases. Glycobiology 16, 29–37 (2006).

61. Lairson, L. L., Henrissat, B., Davies, G. J. & Withers, S. G. Glycosyl transferases: Structures, functions, and mechanisms. Annu Rev Biochem 77, 521–555 (2008).

62. Yicheng Gong. Structure of the priming arabinosyltransferase AftA required for AG biosynthesis of Mycobacterium tuberculosis. Proceedings of the National Academy of Sciences 120, 2017 (2023).

63. Breton, C., Fournel-Gigleux, S. & Palcic, M. M. Recent structures, evolution and mechanisms of glycosyltransferases. Curr Opin Struct Biol 22, 540–549 (2012).

64. Renato Bruni, B. K. High-Throughput Cloning and Expression of Integral Membrane Proteins in Escherichia Coli. Curr Protoc Protein Sci (2014). doi:10.1002/0471140864.ps2906s74.High-throughput.

65. Suloway, C. et al. Automated molecular microscopy: The new Leginon system. J Struct Biol 151, 41–60 (2005).

66. Rizk, S. S. et al. Allosteric control of ligand-binding affinity using engineered conformation-specific effector proteins. Nat Struct Mol Biol 18, 437–444 (2011).

67. Miller, K. R. et al. T cell receptor-like recognition of tumor in vivo by synthetic antibody fragment. PLoS One 7, 1–14 (2012).

68. Dominik, P. K. et al. Conformational Chaperones for Structural Studies of Membrane Proteins Using Antibody Phage Display with Nanodiscs. Structure 24, 300–309 (2016).

69. Kim, J. et al. Structure and drug resistance of the Plasmodium falciparum transporter PfCRT. Nature 576, 315–320 (2019).

70. Punjani, A., Rubinstein, J. L., Fleet, D. J. & Brubaker, M. A. CryoSPARC: Algorithms for rapid unsupervised cryo-EM structure determination. Nat Methods 14, 290–296 (2017).

71. Emsley, P. & Cowtan, K. Coot: Model-building tools for molecular graphics. Acta Crystallogr D Biol Crystallogr 60, 2126–2132 (2004).

72. Slabinski, L. et al. XtalPred: A web server for prediction of protein crystallizability. Bioinformatics 23, 3403–3405 (2007).

73. Adams, P. D. et al. PHENIX: A comprehensive Python-based system for macromolecular structure solution. Acta Crystallogr D Biol Crystallogr 66, 213–221 (2010).

74. Chen, V. B. et al. MolProbity: All-atom structure validation for macromolecular crystallography. Acta Crystallogr D Biol Crystallogr 66, 12–21 (2010).

75. Rouser, G., Fleischer, S. & Yamamoto, A. Two dimensional thin layer chromatographic separation of polar lipids and determination of phospholipids by phosphorus analysis of spots. Lipids 5, 494–496 (1970).

76. Rahlwes, K. C., Puffal, J. & Morita, Y. S. Purification and analysis of mycobacterial phosphatidylinositol mannosides, lipomannan, and lipoarabinomannan. Methods in Molecular Biology 1954, 59–75 (2019).

77. William R. Jacobs Jr., Ganjam V. Kalpana, Jeffrey D. Cirillo, Lisa Pascopella, Scott B. Snapper, Rupa A. Udani, Wilbur Jones, Raúl G. Barletta, B. R. B. Genetic systems for mycobacteria. 204, 537–555 (1991).

78. Hayashi, J. M. et al. Spatially distinct and metabolically active membrane domain in mycobacteria. Proc Natl Acad Sci U S A 113, 5400–5405 (2016).

79. Souza, P. C. T. et al. Martini 3: a general purpose force field for coarse-grained molecular dynamics. Nat Methods 18, 382–388 (2021).

80. Borges-Araújo, L. et al. Martini 3 Coarse-Grained Force Field for Carbohydrates. J Chem Theory Comput 19, 7387–7404 (2023).

81. Brown, C. M. et al. Supramolecular organization and dynamics of mannosylated phosphatidylinositol lipids in the mycobacterial plasma membrane. Proc Natl Acad Sci U S A 120, (2023).

82. Oluwole, A. O. et al. Peptidoglycan biosynthesis is driven by lipid transfer along enzyme-substrate affinity gradients. Nat Commun 13, 1–12 (2022).

83. Alessandri, R. et al. Martini 3 Coarse-Grained Force Field: Small Molecules. Adv Theory Simul 5, (2022).

84. Kroon, P. C. et al. Martinize2 and Vermouth: Unified Framework for Topology Generation. Elife 12, 1–39 (2023).

85. Wassenaar, T. A., Ingólfsson, H. I., Böckmann, R. A., Tieleman, D. P. & Marrink, S. J. Computational lipidomics with insane: A versatile tool for generating custom membranes for molecular simulations. J Chem Theory Comput 11, 2144–2155 (2015).

86. Parrinello, M. & Rahman, A. Polymorphic transitions in single crystals: A new molecular dynamics method. J Appl Phys 52, 7182–7190 (1981).

87. Berendsen, H. J. C., Postma, J. P. M., Van Gunsteren, W. F., Dinola, A. & Haak, J. R. Molecular dynamics with coupling to an external bath. J Chem Phys 81, 3684–3690 (1984).

88. Abraham, M. J. et al. Gromacs: High performance molecular simulations through multi-level parallelism from laptops to supercomputers. SoftwareX 1–2, 19–25 (2015).

89. Lindahl, Abraham, Hess, & van der Spoel. GROMACS 2021.4 Source code. November 5, 2021 | Version 2021.4 https://zenodo.org/records/5636567.

90. Tribello, G. A., Bonomi, M., Branduardi, D., Camilloni, C. & Bussi, G. PLUMED 2: New feathers for an old bird. Comput Phys Commun 185, 604–613 (2014).

91. Hunter, J. D. MATPLOTLIB: A 2D GRAPHICS ENVIRONMENT. Comput Sci Eng 9, 90–95 (2007).

92. Vickery, O. N. & Stansfeld, P. J. CG2AT2: An Enhanced Fragment-Based Approach for Serial Multi-scale Molecular Dynamics Simulations. J Chem Theory Comput 17, 6472–6482 (2021).

93. Lee, J. et al. CHARMM-GUI Membrane Builder for Complex Biological Membrane Simulations with Glycolipids and Lipoglycans. J Chem Theory Comput 15, 775–786 (2019).

94. Gowers, R. et al. MDAnalysis: A Python Package for the Rapid Analysis of Molecular Dynamics Simulations. Proceedings of the 15th Python in Science Conference 98–105 (2016) doi:10.25080/majora-629e541a-00e.

