## Supplementary material for "Mechanistic studies of mycobacterial glycolipid biosynthesis by the mannosyltransferase PimE"

### EXTENDED DATA FIGURE LEGENDS

Extended Data Figure 1

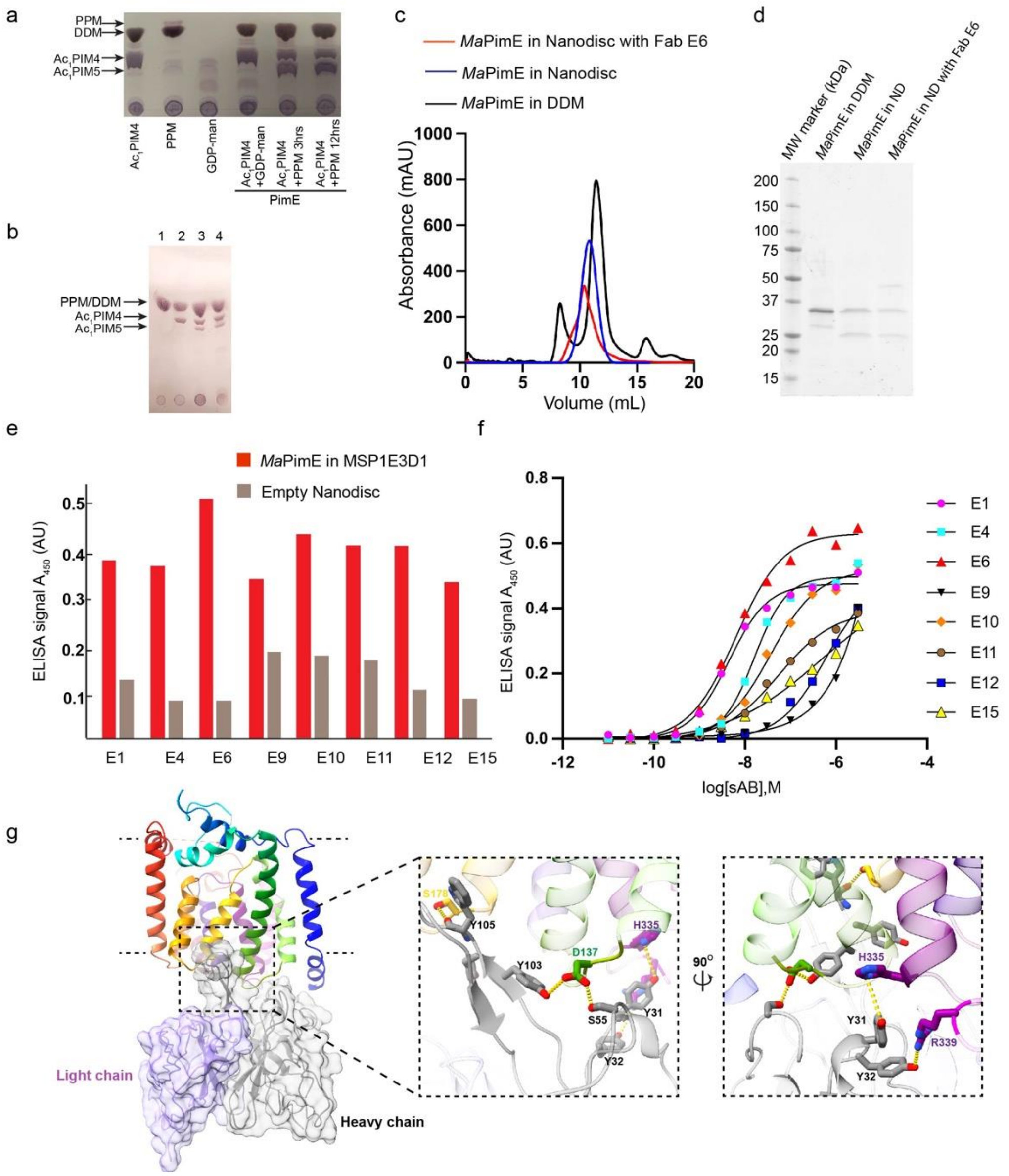


#### Extended Data Fig. 1 Purification and characterization of *Ma*PimE.

(a) TLC analysis of the enzymatic activity of *Ma*PimE in the membrane fraction, confirming the formation of $Ac_1PIM5$ from the substrates $Ac_1PIM4$ and PPM. Glycolipids were visualized by orcinol staining. (b) Metal-independent activity of MaPimE. Lane 1: Membrane fraction with PPM only; Lane 2: Membrane fraction with $Ac_1$PIM4 only; Lane 3: Membrane fraction with both PPM and $Ac_1$PIM4; Lane 4: Membrane fraction with PPM, $Ac_1$PIM4, and metal chelators (EGTA and EDTA). The positions of PPM/DDM, $Ac_1$PIM4, and $Ac_1$PIM5 are indicated by arrows. The presence of Ac1PIM5 in both lanes 3 and 4 indicates that PimE activity is not dependent on metal ions. (c) SEC elution profiles of purified *Ma*PimE in detergent (black), incorporated into a nanodisc (blue) and incorporated into a nanodisc with Fab-E6 bound (red). (d) SDS-PAGE gel of *Ma*PimE purification. The first lane shows *Ma*PimE purified in DDM, the second lane shows *Ma*PimE reconstituted into nanodiscs (using MSP1E3D1 and POPC), and the third lane shows *Ma*PimE reconstituted into nanodiscs (MSP1E3D1 and POPC) with Fab-E6 bound. (e) Single-point ELISA of Fab clones (E1, E4, E6, E9, E10, E11, E12, and E15) against *Ma*PimE, demonstrating the specific binding of the Fabs to *Ma*PimE compared to empty nanodiscs. (f) Multi-point ELISA of selected Fabs against *Ma*PimE, showing the concentration-dependent binding of the Fabs to *Ma*PimE in nanodiscs. (g) The PimE and Fab-E6 binding interface, highlighting the key residues involved in the interaction between the Fab and the cytoplasmic loops of *Ma*PimE.

Extended Data Figure 2

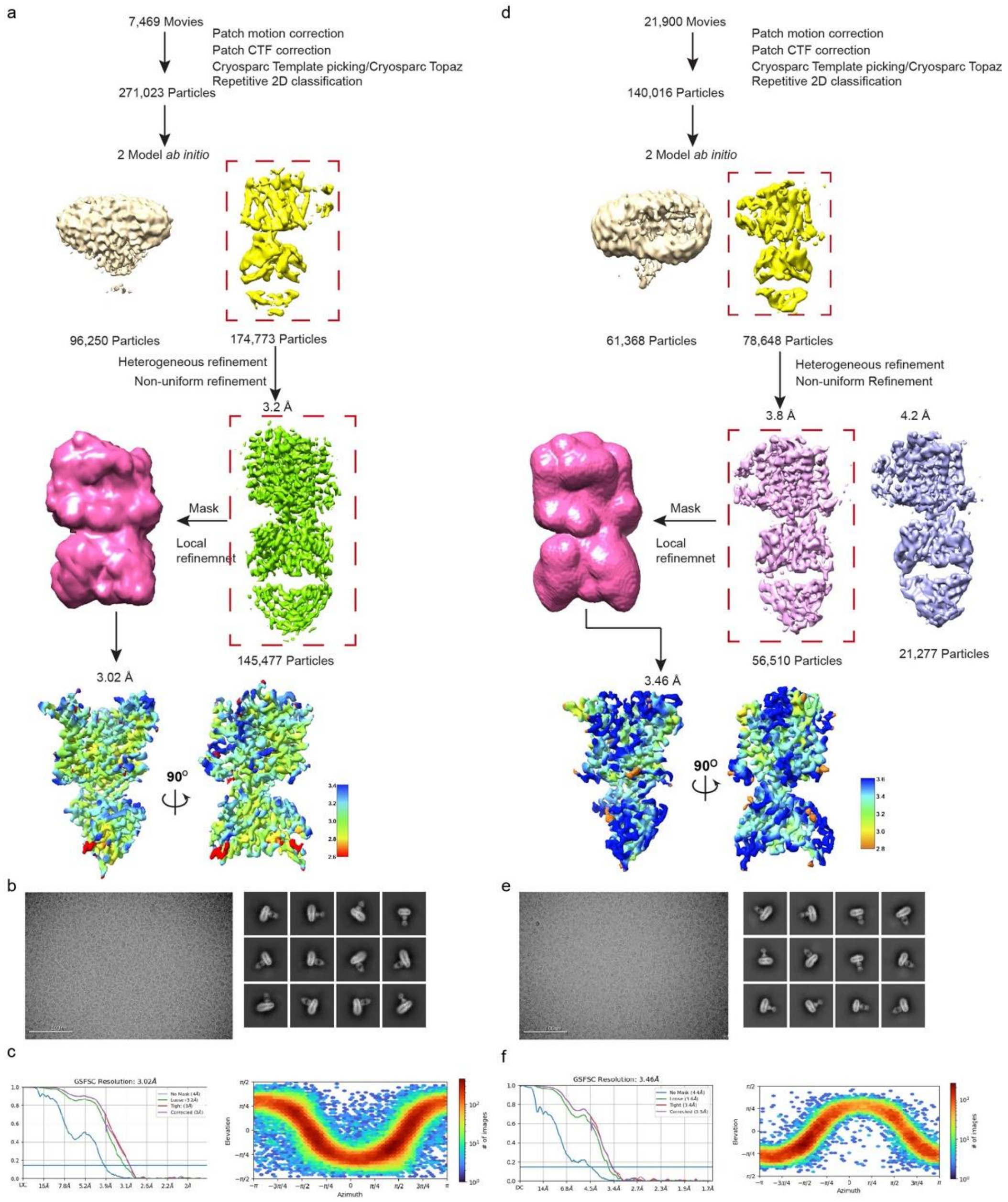


**Extended Data Figure 2. Cryo-EM data processing and validation.**

(a-c) Data processing workflow for the apo structure of *Ma*PimE, including particle picking, 2D classification, *ab initio* model generation, heterogeneous refinement, non-uniform refinement, and local refinement. The final map has a resolution of 3.02 Å (FSC = 0.143). (d-f) Data processing workflow for the substrate-bound structure of *Ma*PimE, following similar steps as in (a-c). The final map has a resolution of 3.46 Å (FSC = 0.143).

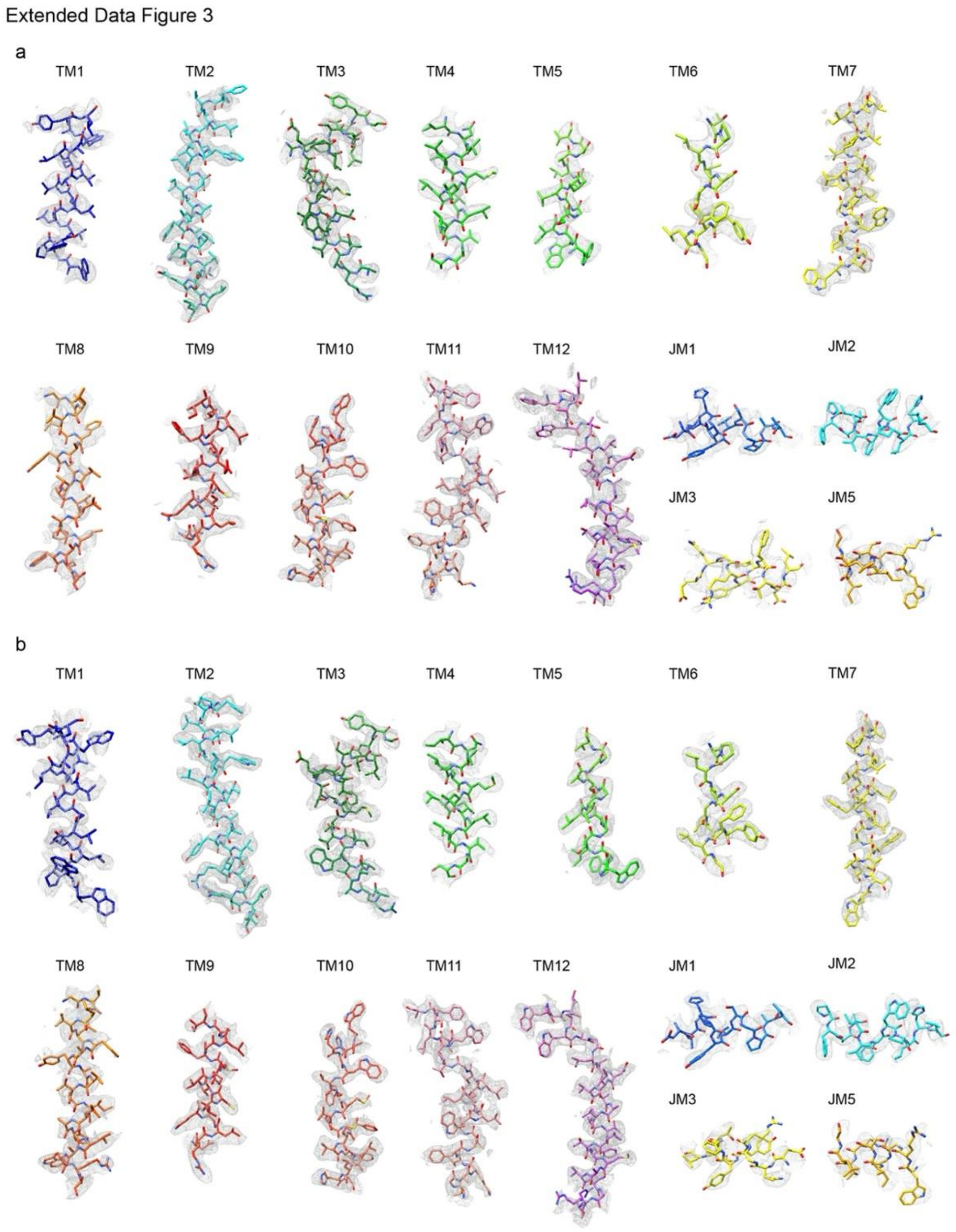


**Extended Data Fig. 3 Cryo-EM density and model for the TM helices and juxtamembrane helices of apo and products-bound *Ma*PimE.**

(a) cryo-EM density (mesh) and atomic model (stick representation) for transmembrane (TM) helices and juxtamembrane (JM) helices in apo *Ma*PimE. (b) cryo-EM density and atomic model for products-bound *Ma*PimE, shown as in (a).

Extended Data Figure 4

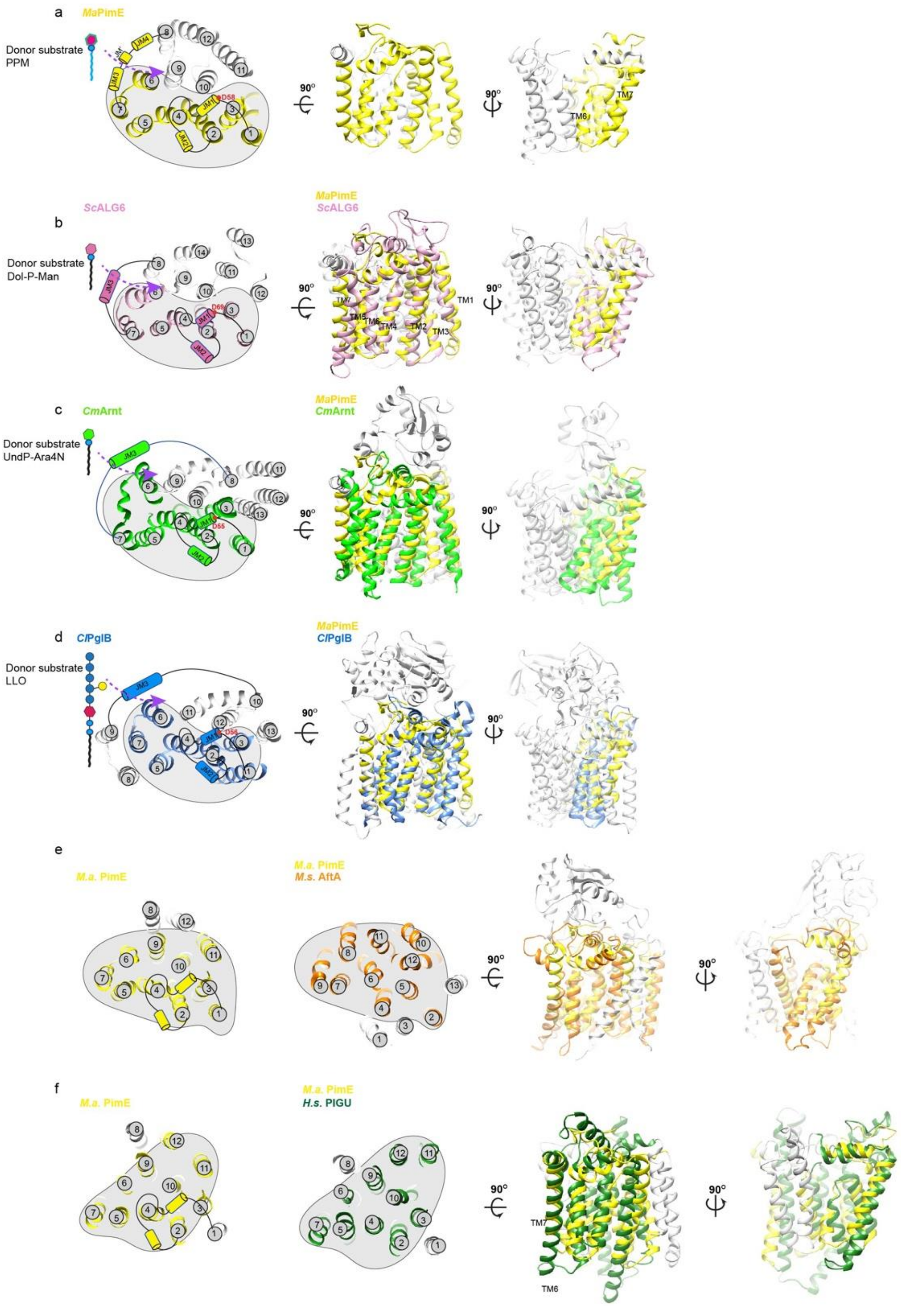

**Extended Data Fig. 4 Comparison of PimE with Structural Homologs.**

(a) The first seven TM helices of PimE are shown in yellow, while the remainder is shown in grey. (b-d) Superposition of PimE with ALG6 (b), ArnT (c), and PglB (d). The first seven TM helices of ALG6, ArnT, and PglB are colored in pink, yellow, and blue, respectively, while the remaining helices are shown in grey. The conserved catalytic residues (D58 in PimE, Asp69 in ALG6, Asp56 in ArnT, and D55 in PglB) are highlighted as sticks. The JM helices (JM1-JM3) and the conserved structural motif formed by the first seven TM helices are labeled. The donor substrates for each enzyme (PPM for PimE, Dol-P-Man for ALG6, UndP-Ara4N for ArnT, and LLO for PglB) are shown as cartoons, revealing a conserved mode of donor substrate binding among these GT-C glycosyltransferases. (e) Superposition of PimE (yellow) with AftA (orange), a mycobacterial arabinosyltransferase, showing structural similarity despite functional differences. The helix bundle of PimE, containing TM helices 1-7 and 9-11, shows a similar architecture to the helix bundle of AftA, which contains TM helices 2 and 4-12. (f) Superposition of PimE (yellow) with PIGU (green), a subunit of the human GPI transamidase complex. The arrangement of the TM helices in PimE, particularly TM helices 2-7 and 9-12, shows a similar architecture to the arrangement of TM helices 2-7 and 9-12 in PIGU.

Extended Data Figure 5

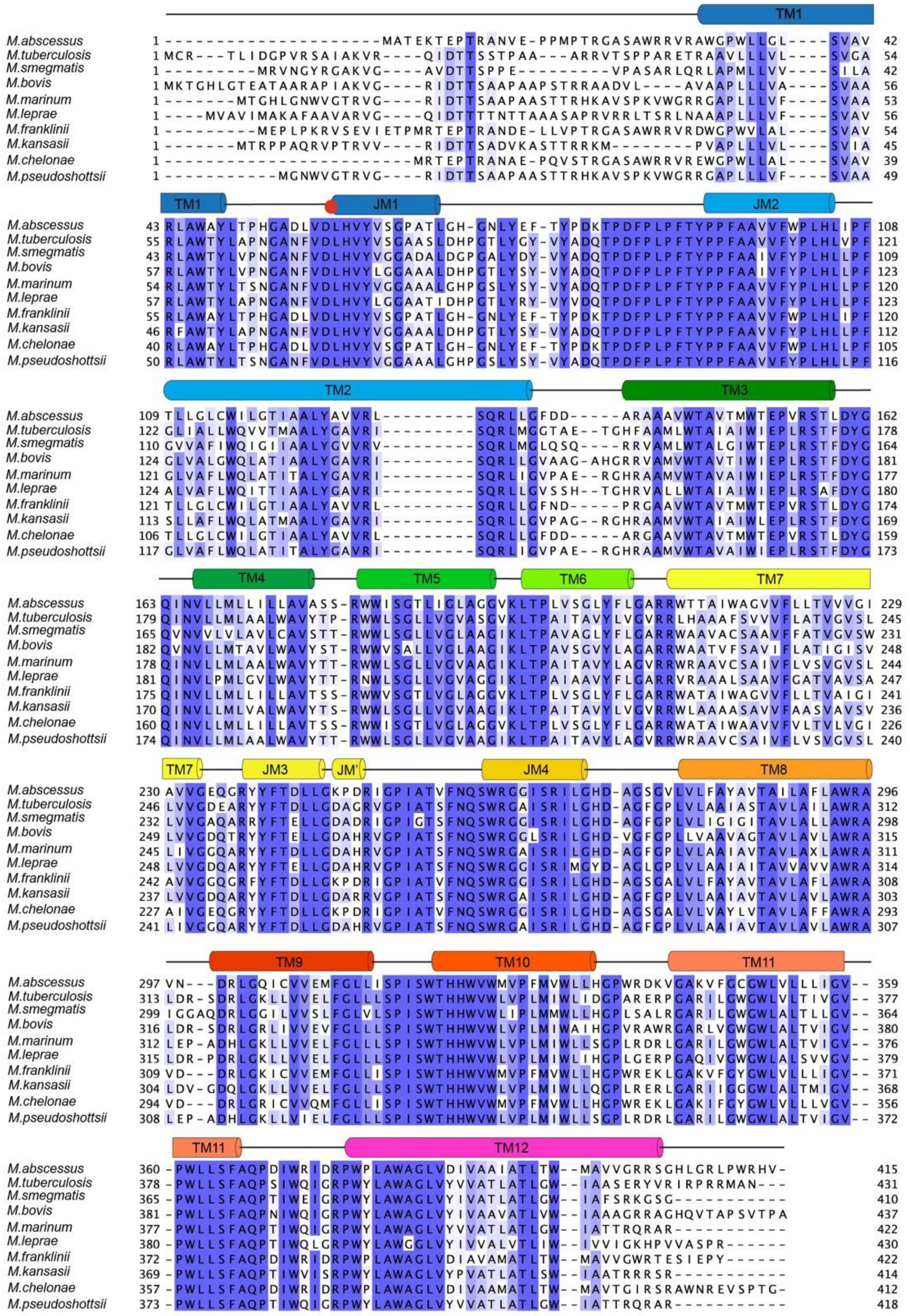

**Extended Data Fig. 5 Sequence alignment of PimE orthologs from various mycobacterial species**. The secondary structure elements (TM helices and JM helices) are indicated and the catalytically essential residue D58 is marked with red dot.

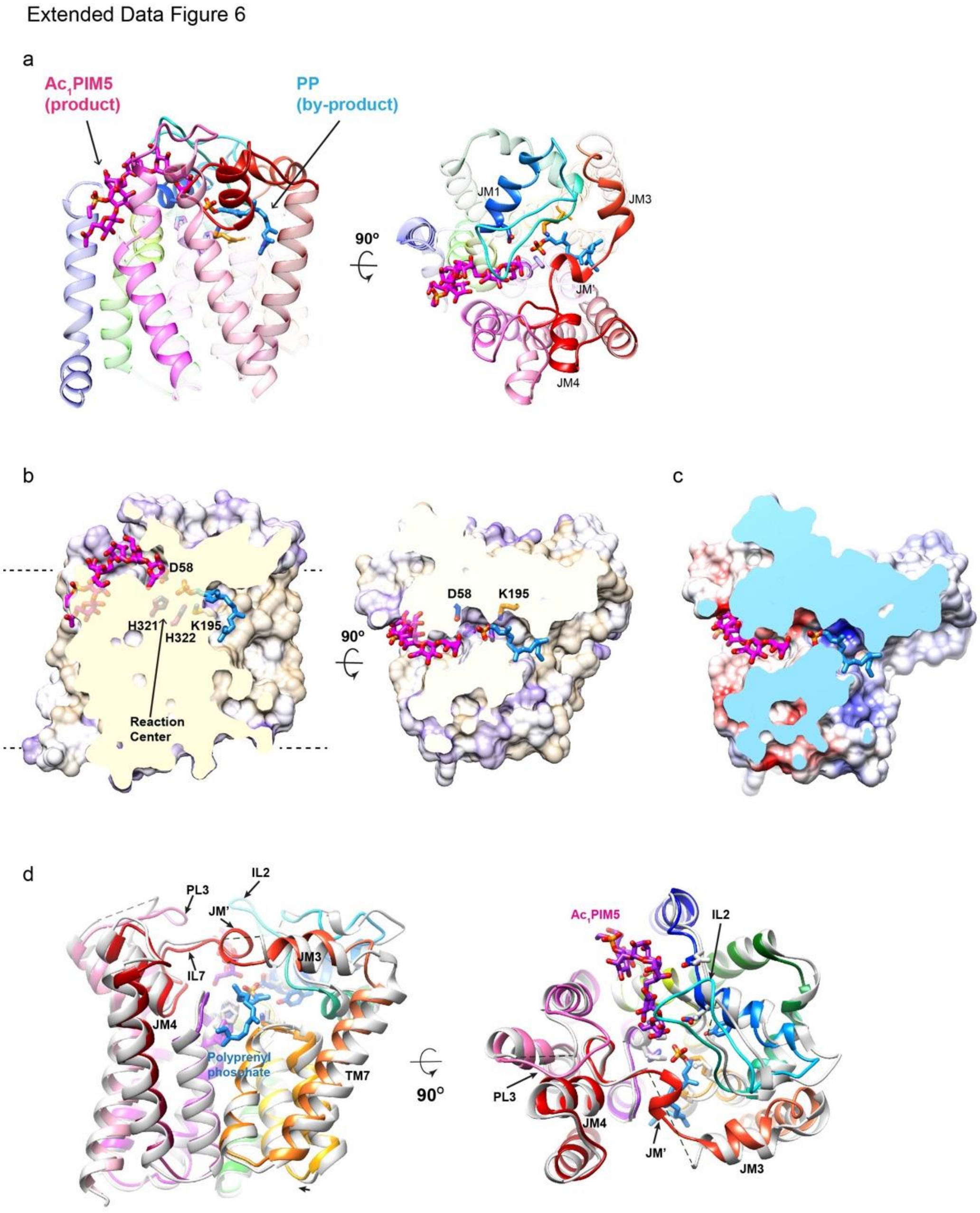


**Extended Data Fig. 6 Product-bound complex of *Ma*PimE.**

(a) Overview of the product-bound structure of *Ma*PimE, focusing on the positioning of PP and *Ac₁PIM5* which are shown as sticks within PimE. (b) Hydrophobic surface representation of the product-bound complex of *Ma*PimE, viewed parallel (left) and perpendicular (right) to the membrane plane. The reaction center, encompassing the conserved residues D58, K195, H321, and H322, is highlighted. The head groups of PP and *$Ac_1$PIM5* converge at the reaction center, with the phosphate group of PP located close to K195 and the fifth mannose of *$Ac_1$PIM5* positioned near D58. (c) Cross-section of the electrostatic surface of the product-bound complex of *Ma*PimE, viewed perpendicular to the membrane plane. *$Ac_1$PIM5* is located along the negatively charged region of the cavity, while PP is situated along the positively charged region. (d) Superimposition of the product-bound *Ma*PimE structure (rainbow) and the apo *Ma*PimE structure (grey) highlighting the conformational rearrangements upon substrate binding, particularly in the periplasmic loops PL1, PL2, and PL3. The TM domain exhibits minimal deviations, with a modest inward pivot rotation/translation of TM helix 7 being the most notable change.

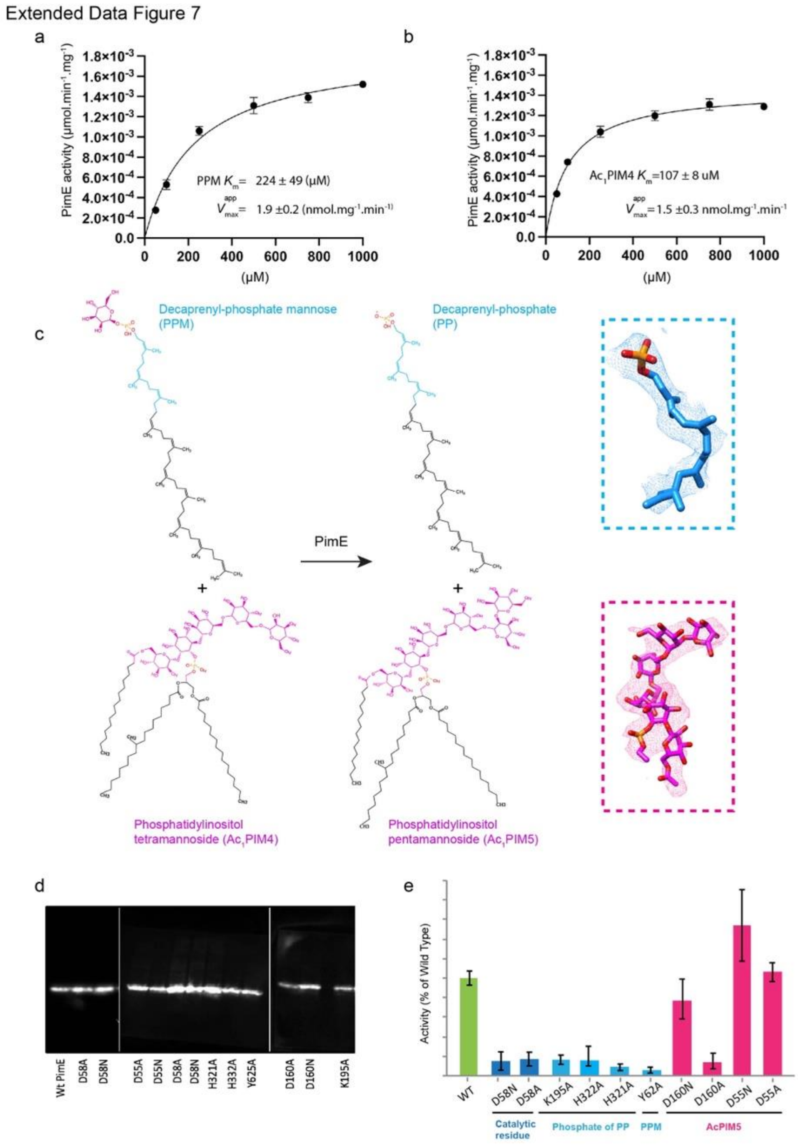


**Extended Data Fig. 7 Kinetic characterization of *Ma*PimE.**

(a-b) Michaelis-Menten plots for PPM (a) and $Ac_1$PIM4 (b), displaying the kinetic parameters ($K_m$ and $V_{max}$) of *Ma*PimE for each substrate. The data points represent the enzymatic activity of *Ma*PimE at different substrate concentrations, with the solid line indicating the best-fit curve according to the Michaelis-Menten equation. The $K_m$ and $V_{max}$ values, along with their standard deviations, are provided for each substrate. (c) Schematic representation of the enzymatic reaction catalyzed by PimE, showing the conversion of $Ac_1$PIM4 and PPM to *$Ac_1$PIM5* and PP. The resolvable density for the carbon chain of PP is colored in blue, while the phosphate group is depicted with heteroatoms (phosphate in orange and oxygen in red). For *$Ac_1$PIM5*, the resolvable density is colored in magenta, with heteroatoms for oxygen (red) and phosphate (orange). (d) Western blot analysis of WT *Ma*PimE and the mutant PimE constructs used for performing the enzymatic assays presented in Fig. 4c. The blot demonstrates that all mutants have expression levels similar to the WT enzyme, ensuring that the observed differences in enzymatic activity can be attributed to the specific mutations rather than variations in protein expression. (e) Relative activity of wild-type and *Ma*PimE mutants expressed in *E. coli*.

Extended Data Figure 8

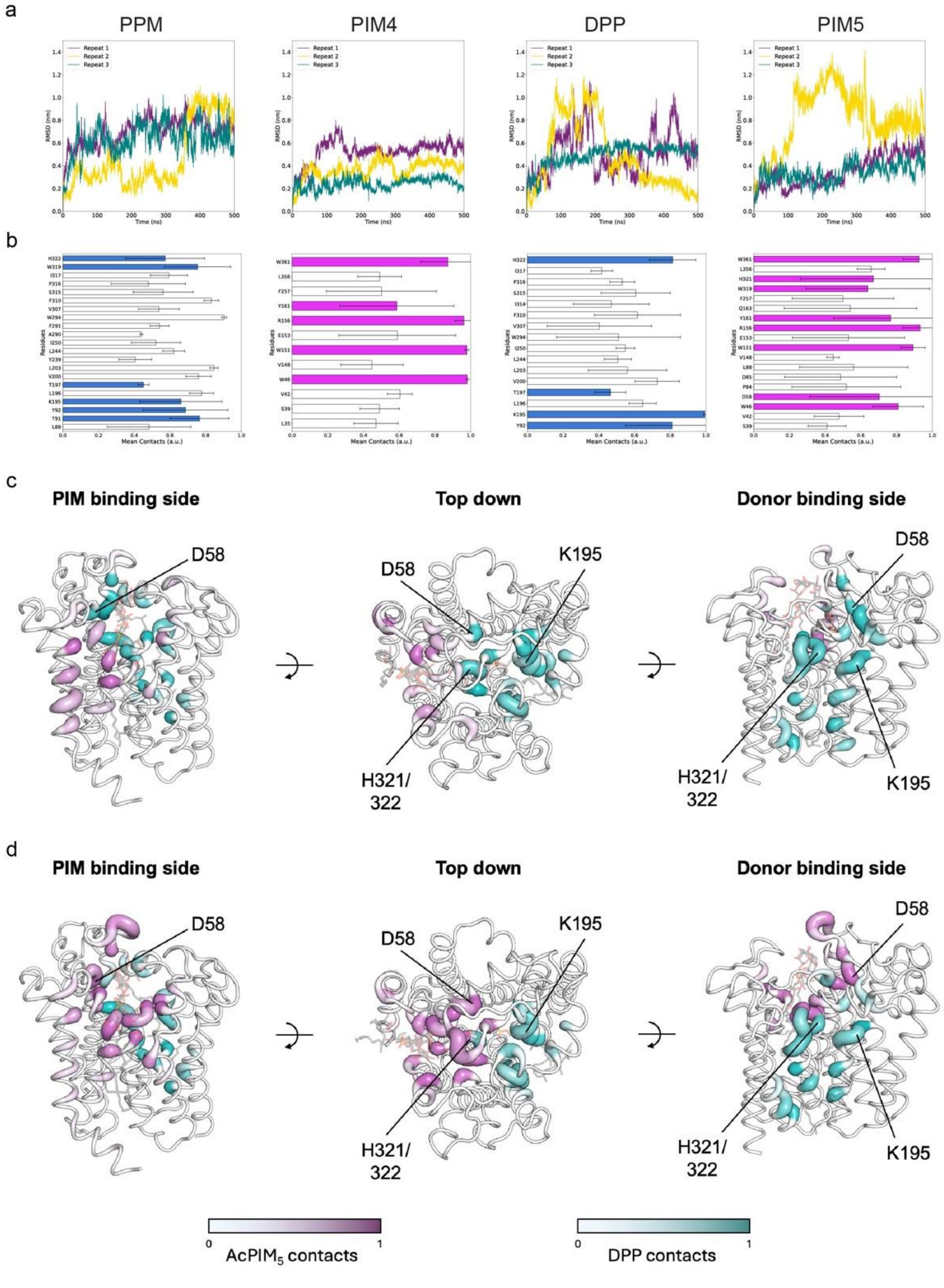


**Extended Data Fig. 8 Molecular dynamics simulations of PimE interactions with substrates and products.**

(a) The root mean squared deviation (RMSD) of the ligands simulated with PimE. The substrates (left) and products (right) are shown, with a line representing an independent repeat. (b) Contact graphs between PimE and ligands over the simulation lengths. The average of three simulations is shown, with the error bar representing standard error. Residues that we subjected to mutation studies are highlighted. A contact value of 1 represents contact with the ligand for the entire simulation; residues with contact values below 0.4 have been omitted for clarity. (c) Areas of PimE in contact with simulated ligands in simulations. The residues in contact with $Ac_1$PIM4 (purple) and PPM (blue). The darker the color, the more contacts throughout all simulations. The cartoon is thicker at regions of higher contact. A contact value of 1 would represent contact with the ligand for the entire simulation. Key residue positions have been highlighted. (d) shows the same as (c), but for $Ac_1$PIM5 (purple) and DPP (blue).

Extended Data Figure 9

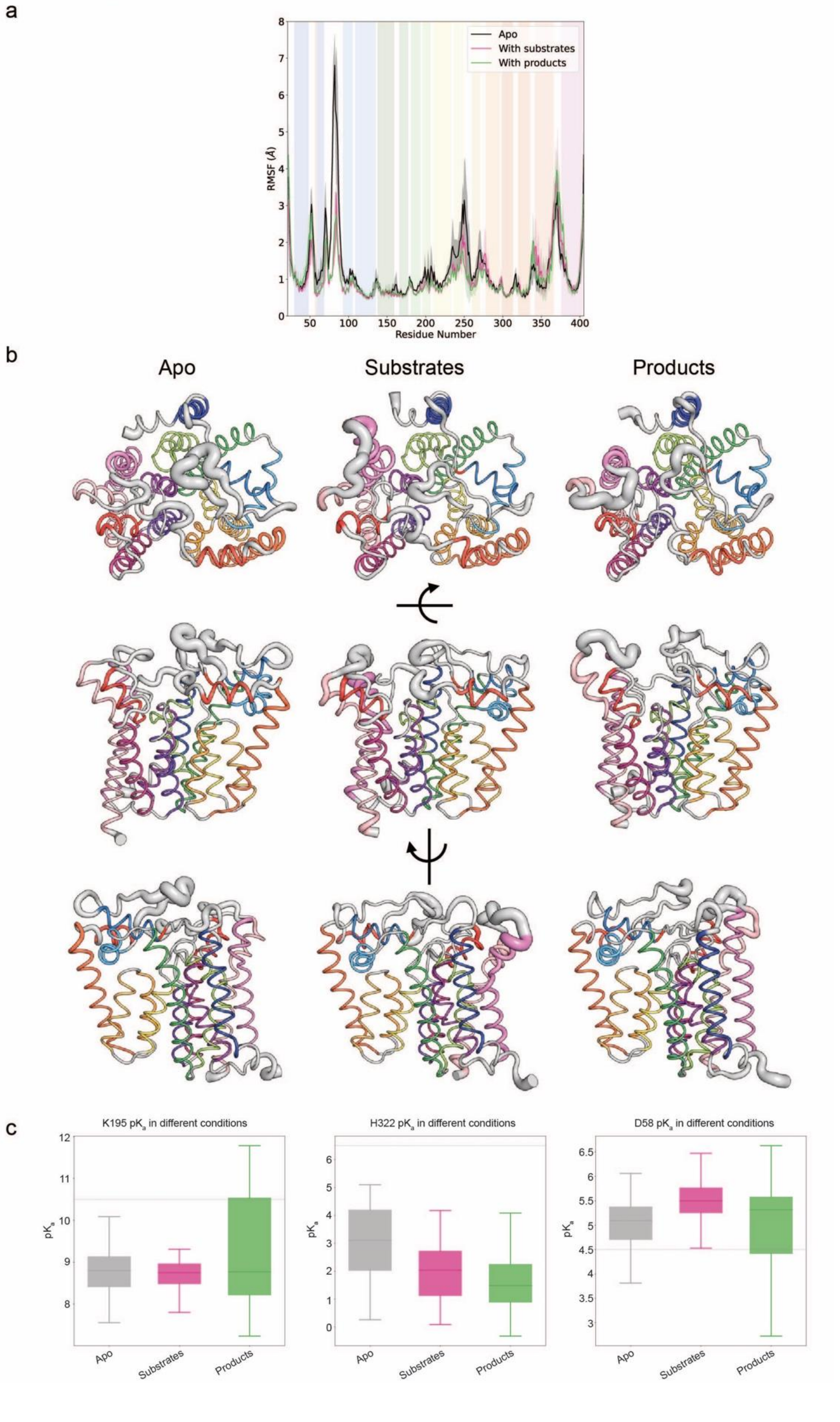

**Extended Data Fig. 9 Molecular dynamics analysis of PimE and protonation states.**

(a) A plot showing the RMSF for each residue as an average of three independent simulations, with the standard error shown as a faint outline. Simulations without substrates shown in black, those with the substrates shown in pink and simulations with products included shown in green. The different regions of the protein as shown in Fig. 1f are highlighted by color. (b) The RMSF shown as cartoon putty on the PimE structures, where the thicker regions are more mobile in simulations. The helices are colored as in Fig. 1f. (c) pKa values of selected residues in different simulation conditions. The gray line represents the expected value for that type of residue. The result is shown for each condition as a summary of all three independent repeats.

Extended Data Figure 10

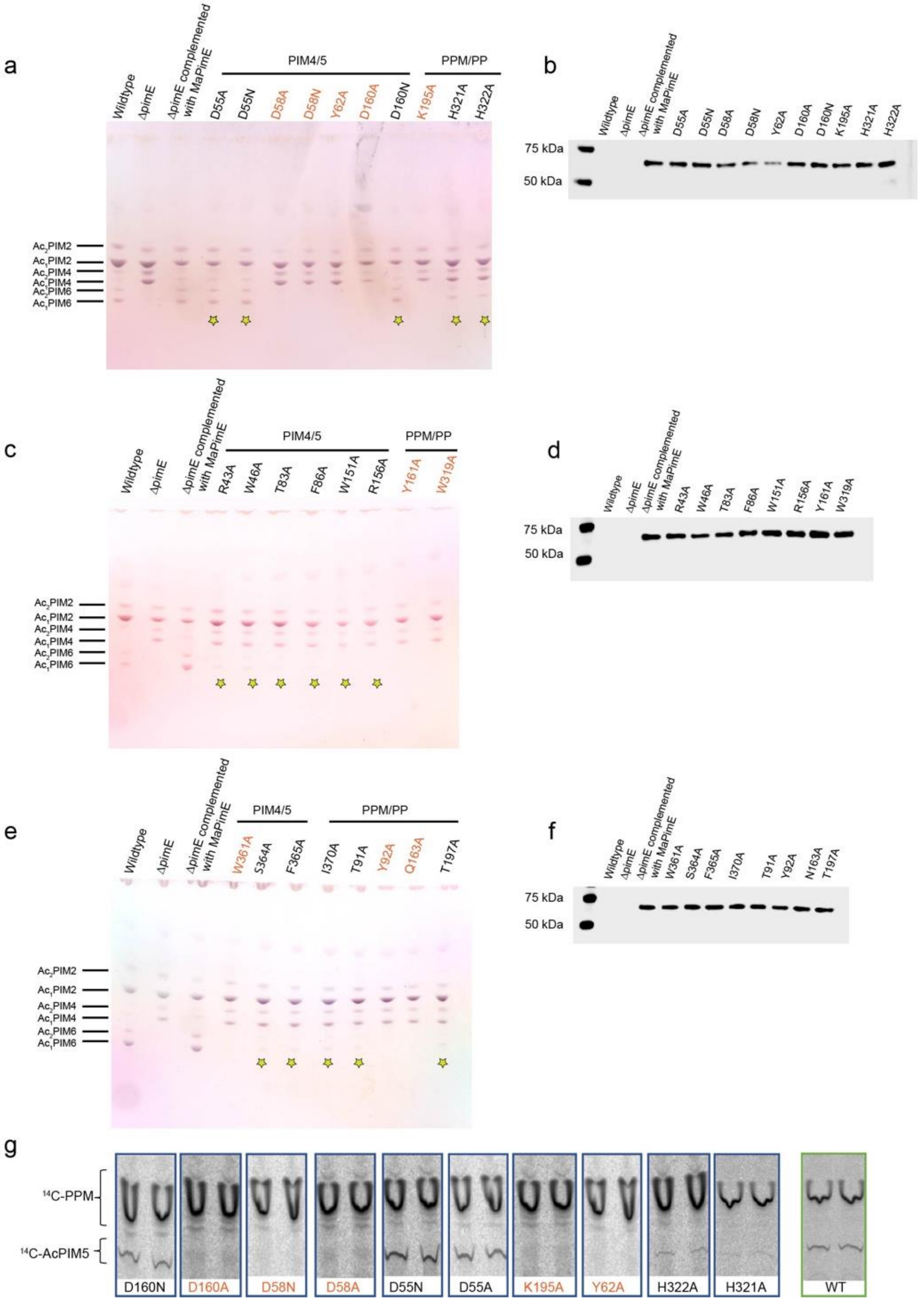


**Extended Data Fig. 10 Functional analysis of PimE mutants in *M. smegmatis.***

(a, c, e) HPTLC analysis of PIMs profiles from *M. smegmatis* strains. Each plate compares PIMs from WT *M. smegmatis* mc²155, Δ*pimE* mutant strain, and Δ*pimE* complemented with WT or mutant *Ma*PimE. Major PIM species are indicated. Yellow stars mark mutants with reduced but not abolished activity (detectable PIM6 production). Mutations causing complete loss of activity are labeled in red-orange. Experiments were performed in triplicate. (b, d, f) Western blot analyses correspond to TLC plates in panels a, c, and e. Blots demonstrate comparable expression levels of WT and mutant *Ma*PimE in complemented strains. (g) *TLC analysis of the enzymatic activity of WT and mutant PimE using* $^{14}$*C-labeled substrates. Mutating D58 to alanine or asparagine abolishes activity, while mutations of K195, H321, H322, and Y62 lead to reduced activity. Mutating D55 does not significantly affect activity, while mutating D160 to alanine largely reduces activity, but mutating it to asparagine only partially reduces activity.*

Table 1 . Cryo-EM data collection and modeling statistics

| | Apo PimE | PimE bound with products |
|---|---|---|
| Data Collection | | |
| Microscope | FEI Titan Krios-CEC | FEI Titan Krios-NYSBC |
| Camera | Gatan K3 | Gatan K3 |
| Voltage (kV) | 300 | 300 |
| Electron expose (e-/Å2) | 58 | 58 |
| Defocus range (μm) | -1.2 to -2.2 | -0.8 to -2.5 |
| Pixel size (Å) | 0.87 | 0.825 |
| Symmetry imposed | C1 | C1 |
| Initial particle images (No.) | 2,047,550 | 10,444,683 |
| Final particle images (No.) | 145,477 | 56,510 |
| Final Resolution (Å) | 3.02 | 3.46 |
| FSC threshold | 0.143 | 0.143 |
| Refinement | | |
| Model composition | | |
| Non-hydrogen atoms | 2882 | 3019 |
| Protein residues | 369 | 384 |
| Ligands | 0 | 2 |
| Waters | 0 | 0 |
| Mean B factor (Å2) | | |
| Protein | 61.7 | 60.7 |
| Ligands | | 43.9 |
| R.m.s. deviation | | |
| Bond lengths (Å) | 0.003 | 0.003 |
| Bond angles (°) Validation | 0.46 | 0.48 |
| Clashscore | 4 | 8 |
| Rotamers outlier (%) | 0 | 0 |
| Ramachandran plot | | |
| Favored (%) | 96 | 95 |
| Allowed (%) | 4 | 5 |
| Disallowed (%) | 0 | 0 |

**Table 1.** Cryo-EM data collection and modeling statistics for the apo and substrate-bound structures of *Ma*PimE.
